# Apoptotic Extracellular Vesicles (ApoEVs) Safeguard Liver Homeostasis and Regeneration *via* Assembling an ApoEV-Golgi Organelle

**DOI:** 10.1101/2021.02.24.432630

**Authors:** Bingdong Sui, Runci Wang, Chider Chen, Xiaoxing Kou, Di Wu, Ho-Chou Tu, Yanzhuang Wang, Yijing Liu, Orit Jacobson, Xiaoyuan Chen, Haixiang Liu, Ryan Tsz Kin Kwok, Ben Zhong Tang, Hexin Yan, Minjun Wang, Lei Xiang, Xutong Yan, Yu Fu, Xiao Zhang, Jianxia Tang, Lan Ma, Lu Lu, Yan Jin, Songtao Shi

## Abstract

Apoptosis is an integral physiological cell death process that occurs frequently and generates a huge number of apoptotic extracellular vesicles (apoEVs). However, whether apoEVs are necessary for maintaining organ homeostasis remains unclear. Here, we show that circulatory apoEVs engraft in liver and undergo specialized internalization by hepatocytes (HCs) based on surface signature of galactose and *N*-acetylgalactosamine. Furthermore, apoEVs rescue liver injury in apoptotic-deficient *Fas* mutant and *Caspase-3* knockout mice, which is exerted by restoring the featured hepatic ploidy homeostasis. Surprisingly, apoEVs form a chimeric organelle complex with recipient Golgi apparatus *via* SNARE-mediated membrane interaction, which consequently facilitates microtubule organization and HC cytokinesis. Notably, through Golgi recovery and ploidy transition, apoEVs contribute to liver regeneration and protect against acute hepatic failure. Collectively, these results identify a previously unrecognized role for apoEVs and the specific mechanisms by which they safeguard liver homeostasis, and suggest the potential of apoEV-based therapy for liver disorders.

## Introduction

Apoptosis is a form of programmed cell death that primarily accounts for routine cell turnover in healthy individuals; it is estimated to occur in the human body at a rate of approximately 150 billion cells or roughly 0.4% of total cells per day (Arandjelovic and Ravichandran, 2015; Elliott and Ravichandran, 2016; Morioka et al., 2019). Such physiological cell death is recognized as an integral part of development and homeostatic maintenance, while it also contributes to tissue stress responses and regeneration (Fuchs et al., 2013; Fuchs and Steller, 2011). During apoptosis, cells undergo a highly regulated process of disassembly including membrane blebbing and protrusion, which ultimately leads to generation of apoptotic fragments as extracellular vesicles (EVs) (Atkin-Smith et al., 2015; Coleman et al., 2001; Poon et al., 2014). The heterogeneity of these EVs, originally known as apoptotic bodies (apoBDs), has recently begun to be appreciated (Caruso and Poon, 2018). They vary in their mechanisms of production, cargos, and size ranging from 50 nm to several microns; for these reasons, the more general term of apoptotic extracellular vesicles (apoEVs) is now favored (Caruso and Poon, 2018; Dieude et al., 2015; Lynch et al., 2017). Tens of apoEVs can be released per cell, and they have increasingly been noticed as regulators of immune responses, stem cell-mediated tissue turnover and regenerative therapies (Berda-Haddad et al., 2011; Brock et al., 2019; Liu et al., 2018; Liu et al., 2020). However, the essential role of apoEVs in maintaining organ homeostasis remains underdetermined.

The liver has long been considered as a major organ for apoptotic cell removal from circulation, a function which is mostly attributed to the resident macrophages, called Kup□er cells (Shi et al., 1996). It is generally believed that efficient phagocytosis is tolerogenic and critically maintains liver homeostasis, whereas defective efferocytosis contributes to a spectrum of hepatic pathologies due to overwhelming apoptotic material-induced hyperinflammation (Morioka et al., 2019). Engulfment of apoEVs by liver macrophages can nevertheless be pro-inflammatory and fibrogenic (Canbay et al., 2003), leaving the detailed roles played by apoEVs in the liver controversial and elusive. Notably, the liver is also well known for its extreme and rapid capacity to regenerate after injuries, while loss or impairment of this regenerative capacity underlies severe or chronic liver damage that may eventually progress to hepatic failure (Forbes and Newsome, 2016). It has been reported that mice lacking either caspase-3 or caspase-7, the key apoptotic executors, demonstrate a substantial defect in liver regeneration after partial hepatectomy (PHx) (Li et al., 2010). Delayed liver regeneration after PHx has also been documented in Fas-deficient lymphoproliferative (*Fas^lpr^*) mice, which show defective Fas-induced apoptosis and resemble human systemic lupus erythematosus, an autoimmune disease associated with increased susceptibility to liver disease (Bessone et al., 2014; Desbarats and Newell, 2000; Watanabe-Fukunaga et al., 1992). In light of these findings, understanding the regulatory function and mechanisms of apoEVs in the liver will have significant implications for deciphering principles that underlie hepatic regeneration and establishing novel therapies for hepatic disease.

EVs have emerged as playing pivotal roles in intercellular communication based on their surface bioactive molecules and various types of cargos including nucleic acids, proteins and lipids (Pitt et al., 2016; Roy et al., 2018). Given these properties, EVs serve as homeostatic modulators and can be leveraged in both diagnostics and therapeutics (Pitt et al., 2016; Roy et al., 2018). ApoEVs, however, are a relatively understudied EV population which possess unique molecular signatures (Dieude et al., 2015; Wickman et al., 2013). Despite the characterization of EVs and their potent functions, how these vesicles recognize recipient cells and what processes they undergo in recipient cells are not yet fully understood. The current model postulates uptake of EVs *via* endocytosis, after which they shuttle within endosomes, surf on the endoplasmic reticulum and finally fuse with acidified endosomes or lysosomes for cargo release (Heusermann et al., 2016; Joshi et al., 2020). Nevertheless, we have limited knowledge of the specific mechanisms that EVs use to target certain recipient cells and the biological effects that EVs may exert on the recipient endomembrane system.

Here, we aimed to investigate the biological properties and related physiological function of apoEVs in maintenance of organ homeostasis. We discovered that apoEVs are characterized by specific surface markers which endow them with remarkable liver engraftment, specialized internalization by hepatocytes (HCs) and functional assembly with recipient Golgi apparatus. ApoEVs were further revealed to crucially preserve hepatic ploidy homeostasis, maintain liver integrity, promote liver regeneration after PHx and mediate therapy against acute liver failure (ALF) in mice. These findings suggest a general understanding of the characterization, function, mechanisms and application of apoEVs underlying organ maintenance and regeneration, with potential applications for novel therapeutics.

## Results

### Circulatory apoEVs engraft in the liver and safeguard liver homeostasis

ApoEVs used in this study were induced from multiple cell types (stem/stromal cells, immune cells and fibroblasts) and different species (human and mouse) and by various methods (protein kinase inhibition, oxidative stress, mitochondrial permeabilization and Fas crosslinking) targeting both the intrinsic (*i.e.* mitochondrial) and extrinsic (*i.e.* death receptor) pathways of apoptosis (Xu and Shi, 2007). Among these, the apoEVs primarily discussed here are those derived from mesenchymal stem cells (MSCs) after staurosporine (STS) treatment (Liu et al., 2018; Liu et al., 2020). Notably, the protein kinase inhibitor STS has been recognized as an apoptosis inducer *via* both caspase-dependent and caspase-independent pathways (Belmokhtar et al., 2001). Moreover, MSCs represent potent EV donors with immense translational potential, and require the apoptotic process to fulfill their therapeutic effects after transplantation (Galleu et al., 2017; Liu et al., 2020). The protocol was determined according to our previously established methodology for isolating apoBDs (Liu et al., 2018; Liu et al., 2020) with necessary modifications for collecting the general apoEV population more broadly (Figure S1A). STS-induced apoptosis of MSCs was confirmed by morphology (Figure S1B) and terminal deoxynucleotidyl transferase dUTP nick end labeling (TUNEL) analysis (Figure S1C). Notably, the protein yield of collected apoEVs was 10-fold higher than that of exosomes (Exos) released by MSCs in the non-apoptotic culture condition (Figure S1D). ApoEVs demonstrated characteristic morphology (Figure S1E) with a wide size distribution between 100-800 nm in diameter (Figure S1F). Further analyses revealed that the apoEVs expressed apoptotic markers such as cleaved caspase-3 and surface exposures of typical phagocytosing signals (Arandjelovic and Ravichandran, 2015) phosphatidylserine (PtdSer, shown by Annexin V binding), complement component 1q (C1q) and thrombospondin 1 (TSP1) (Figure S1G-J). Nevertheless, apoEVs did not express the nuclear lamina component Lamin B1, a negative marker of EVs (Figure S1J). Therefore, apoEVs were verified with their specialized profiles being discovered.

To investigate the potential physiological function of apoEVs, we first examined the biodistribution of circulatory apoEVs by tracing labeled apoEVs after intravenous infusion. Positron emission tomography (PET) scanning of live mice demonstrated that copper radioisotope (^64^Cu)-labeled apoEVs engraft in liver, among other tissues, as early as 1 h after infusion (Figure 1A). Surprisingly, in a 48 hours period, the ^64^Cu radioisotope signal indicating apoEVs increasingly aggregated in liver, with approximately 70% of total ^64^Cu-apoEV signal detected in liver at 48 h; at this point, the liver had 10-fold higher ^64^Cu-apoEV radiation intensity than any other dissected organ/tissue (Figure 1A). Tracing of apoEVs labeled with the lipophilic fluorescent dye PKH26 confirmed that liver engraftment after infusion proceeded in a manner resembling zero-order elimination kinetics, *i.e.* a constant amount being eliminated per unit time, with a half-time (*t*_1/2_) of about 8 days (Figure 1B). The linear change indicated that the liver biologically processed or metabolized apoEVs.

**Figure 1.**
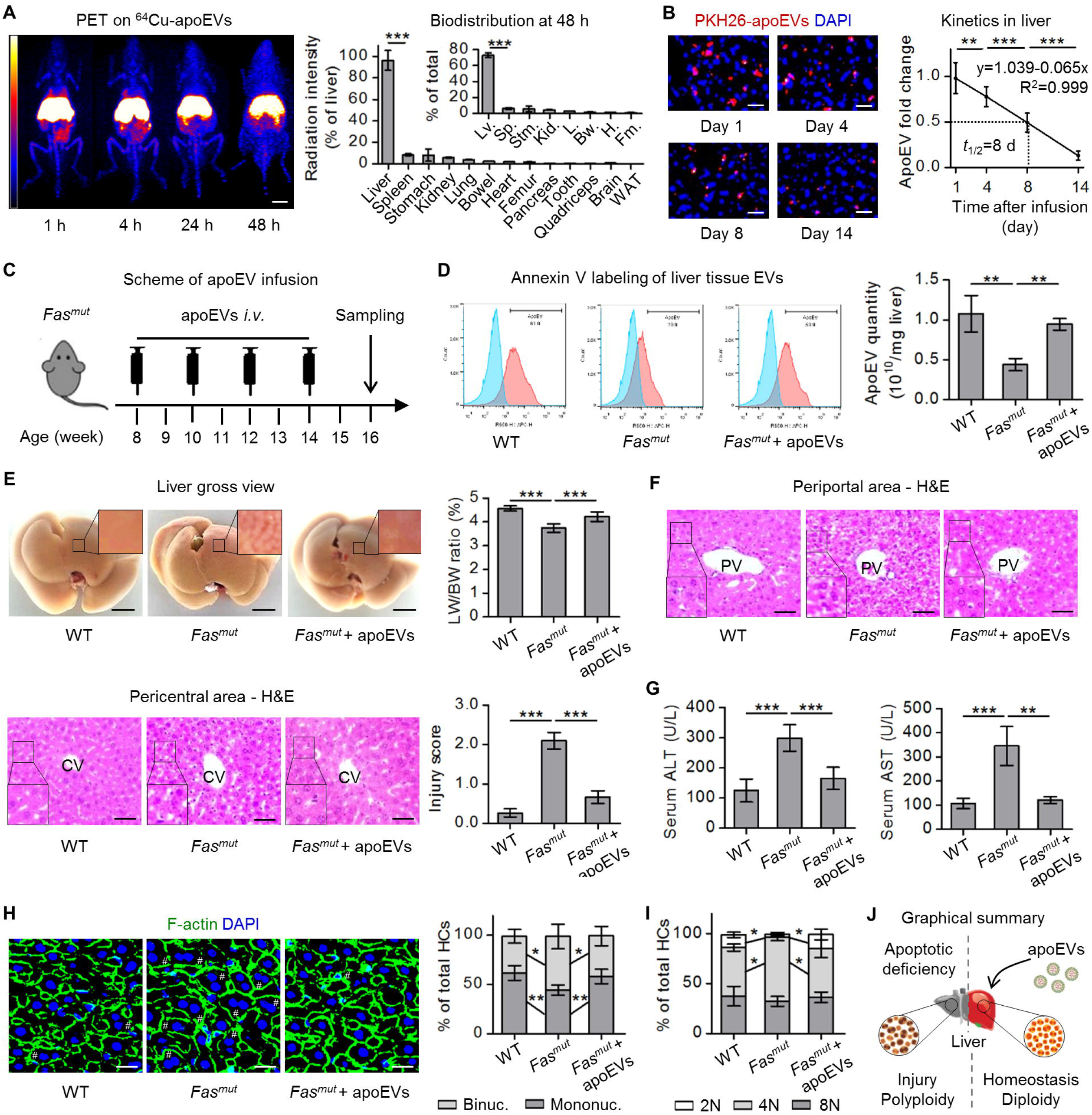
Circulatory apoptotic extracellular vesicles (apoEVs) engraft in the liver and safeguard liver homeostasis. **(A)** Biodistribution of intravenously injected copper isotope (^64^Cu)-labeled apoEVs. Mice underwent whole-body coronal positron emission tomography (PET) at indicated time points. Organs/tissues were dissected and measured for radiation intensity at 48 h. WAT, white adipose tissue; Lv., liver; Sp., spleen; Stm., stomach; Kid., kidney; L., lung; Bw., bowel; H., heart; Fm., femur. Scale bar = 10 mm. *N* = 4 per organ/tissue. **(B)** Tracing of PKH26-labeled apoEVs (red) in the liver, counterstained with DAPI for detecting DNA (blue). ApoEVs were intravenously injected and liver samples were collected at indicated time points. After a single infusion, kinetic changes of apoEVs in the liver were obtained by fluorescent imaging analysis. The half-life of liver-engrafted apoEVs was calculated. Scale bars = 25 μm. *N* = 6 per time point. **(C)** Schematic diagram showing study design of apoEV infusion in *Fas* mutant (*Fas^mut^*) mice. ApoEVs were infused intravenously (*i.v.*) at a protein concentration of 5 μg/g. **(D)** Flow cytometric analysis of Annexin V^+^ liver tissue extracellular vesicles (apoEVs) with corresponding quantification. WT, wild type. *N* = 3 per group. **(E)** Gross view images of liver and quantification of fasting liver weight (LW) to body weight (BW) ratio. Scale bars = 5 mm. *N* = 6 per group. **(F)** Hematoxylin and eosin (H&E) staining of liver tissues in respective periportal vein (PV) and pericentral vein (CV) areas. Hepatic injury scores were examined based on pathological parameters. Scale bars = 50 μm. *N* = 6 per group. **(G)** Serum alanine aminotransferase (ALT) and aspartate aminotransferase (AST) levels were determined. *N* = 6 per group. **(H)** Representative liver fluorescent images showing hepatocytes (HCs) with different numbers of nuclei (blue, DAPI for DNA) and their cell borders (green, phalloidin for F-actin). # indicates binucleated HCs. Percentages of binucleated and mononucleated HCs were quantified. Scale bars = 25 μm. *N* = 6 per group. **(I)** After propidium iodide (PI) staining, ploidy distribution of HCs (diploid, 2N; tetraploid, 4N; octoploid, 8N) was analyzed by flow cytometry. *N* = 3 per group. **(J)** Graphical summary demonstrating that apoEVs maintained liver homeostasis in apoptotic deficient conditions. Data represent mean ± standard deviation. *, *P* < 0.05; **, *P* < 0.01; ***, *P* < 0.0001. Statistical analyses were performed by one-way analysis of variance followed by the Newman-Keuls post hoc tests.

The above data inspired us to explore whether hepatic apoEVs were indeed associated with any functional implications. To address this issue, we studied two mouse models with deficiency in apoptosis (Liu et al., 2018), specifically *Fas* mutant (*Fas^mut^*, characterized by functionally deficient Fas, previously referred to as the *Fas^lpr^* mice (Watanabe-Fukunaga et al., 1992)) and *Caspase-3* knockout (*Casp3^-/-^*) mice. As has been previously established (Xu and Shi, 2007), the Fas ligand (FasL)/Fas pathway is a well-characterized extrinsic apoptosis pathway, while caspase-3 is critical for the final execution phase of both intrinsic and extrinsic pathways of apoptosis. We confirmed that *Fas^mut^* and *Casp3^-/-^* mice are both defective in physiological apoptosis, as shown by decreased circulatory and organ-specific cellular apoptotic rates in freshly isolated peripheral blood mononucleated cells (PBMNCs) (Figure S1K), bone marrow mononucleated cells (BMMNCs) (Figure S1L) and liver non-parenchymal cells (NPCs) (Figure S1M). We have also introduced a liver EV isolation protocol (Figure S1N) (Ishiguro et al., 2019) and applied Annexin V binding to detect a tissue-specific apoEV population (Liu et al., 2018). The liver EVs were identified based on morphology, particle distribution, expression of tetraspanins CD9, CD63 and CD81, and negative for Lamin B1 (Figure S1O-Q).

A chronic intermittent apoEV infusion protocol at a biweekly interval was designed based on the hepatic elimination kinetics to replenish hepatic apoEVs (Figure 1C). Notably, systemic infusion of apoEVs had limited influence on general traits of *Fas^mut^* mice in terms of body weight (BW) gain (Figure S2A), food intake (Figure S2B) and daily spontaneous activity (Figure S2C). Also notable is that despite being recognized as a lymphoproliferative model (Watanabe-Fukunaga et al., 1992), *Fas^mut^* mice used in this study (8-16 weeks of age on the B6 background) did not show inflammatory features and their inflammation level remained comparable to wild type (WT) upon apoEV infusion (Figure S2D). As expected, we discovered that *Fas^mut^* mice possessed less than half amount of liver apoEVs as WT mice, a deficiency which was recovered by apoEV infusion (Figure 1D). The total liver tissue EV quantity, however, was unchanged despite *Fas^mut^* and apoEV infusion (Figure S2E), indicating the ability of the organism to adapt and maintain the tissue EV pool at a constant level. Importantly, partial loss of apoEVs in the liver of *Fas^mut^* mice was functionally sufficient to provoke liver injury, as demonstrated by a reduced fasting liver weight (LW) to BW ratio, obvious alterations upon gross examination and in histological analysis, and elevated serum levels of alanine aminotransferase (ALT) and aspartate aminotransferase (AST) (Figure 1E-G). ApoEV infusion restored the liver tissue integrity in *Fas^mut^* mice (Figure 1E-G) without triggering fibrotic or inflammatory responses (Figure S2F, G). Furthermore, apoEV infusion also rescued the liver damage developed by *Casp3^-/-^* mice due to hepatic apoEV deficiency (Figure S3A-E).

In analyzing liver tissue injuries, we noticed with interest that *Fas^mut^* mice tended to possess a high percentage of binucleated HCs, and that cytoplasm vacuolization occurred in a high prevalence in these binucleated HCs (Figure 1F). One cellular characteristics of the mammalian liver is the polyploidization of HCs, during which individual HCs acquire more than two sets of chromosomes resulting from the failure of cytokinesis (Wang et al., 2017). Progressive polyploidization is considered a pathophysiological process in the liver (Gentric et al., 2015), but the detailed contributions of polyploid HCs to liver function and their regulatory mechanisms are not fully understood (Wilkinson et al., 2019a; Wilkinson et al., 2019b; Zhang et al., 2018a). We further evaluated whether apoEVs regulate the HC ploidy status in the liver. Phalloidin staining revealed HC cell borders and demonstrated that *Fas^mut^* mice aggregated more binucleated HCs than mononucleated HCs compared to WT mice, which was rescued by apoEV infusion (Figure 1H). Moreover, propidium iodide (PI) staining of HC chromatin showed that *Fas^mut^* mice possessed reduced diploid (2N) HCs and increased tetraploid (4N) HCs compared to WT mice, which was also rectified by apoEV infusion (Figure 1I). Octoploid (8N) HCs were not significantly influenced in the experimental context. ApoEV infusion also restored the ploidy homeostasis in *Casp3^-/-^* mouse livers, which developed polyploidization as well (Figure S3F, G). Taken together, these data suggest that apoEVs are indispensably required for safeguarding liver ploidy and functional homeostasis (Figure 1J).

### Specialized internalization of apoEVs by HCs promotes ploidy transition in the liver

Next, we investigated how apoEVs regulate liver homeostasis. Considering the clues that apoEVs may contribute to the reversal of HC polyploidization, we examined whether HCs, particularly binucleated ones, uptake apoEVs directly for their functional regulation. By using PKH26 to label the membrane of apoEVs before infusion, we confirmed that binucleated HCs in the liver were indeed capable of taking up apoEVs, such that PKH26 fluorescent signals were detected in approximately 40% of HCs at 24 hours after infusion (Figure 2A). PKH26-labeled infused apoEVs were also engulfed by F4/80^+^ liver-resident macrophages (Figure S2H). Given that lipophilic membrane labeling may be xenobiotic and result in loss of membrane integrity, among other drawbacks, we further applied a previously established mitochondria-targeting luminogen, known as DCPy, with aggregation-induced emission (AIE) characteristics (Zheng et al., 2020). This AIE luminogen (AIEgen) is a photosensitizer which generates singlet oxygen (^1^O_2_) upon light irradiation and induces apoptosis (Zheng et al., 2020; Zheng et al., 2018). We identified that DCPy e□ciently labeled apoEVs after apoptotic induction, and that the AIEgen-labeled apoEVs were mainly internalized by HCs after the infusion (Figure 2A, S2H), suggesting that HCs are the primary cellular targets of apoEVs in the liver. To further examine whether the HC uptake of apoEVs occurred as a physiological event, we applied a parabiosis model in which *Fas^mut^* apoptosis-deficient mice were connected with green fluorescent protein (GFP) transgenic mice (*GFP^+/+^*) to obtain shared circulatory apoEVs (Liu et al., 2018). Indeed, GFP signals were detected in binucleated HCs of the *Fas^mut^* mice (Figure 2B). These data indicated a putative link between internalization of apoEVs by HCs, particularly binucleated ones, and functional regulation of the liver.

**Figure 2.**
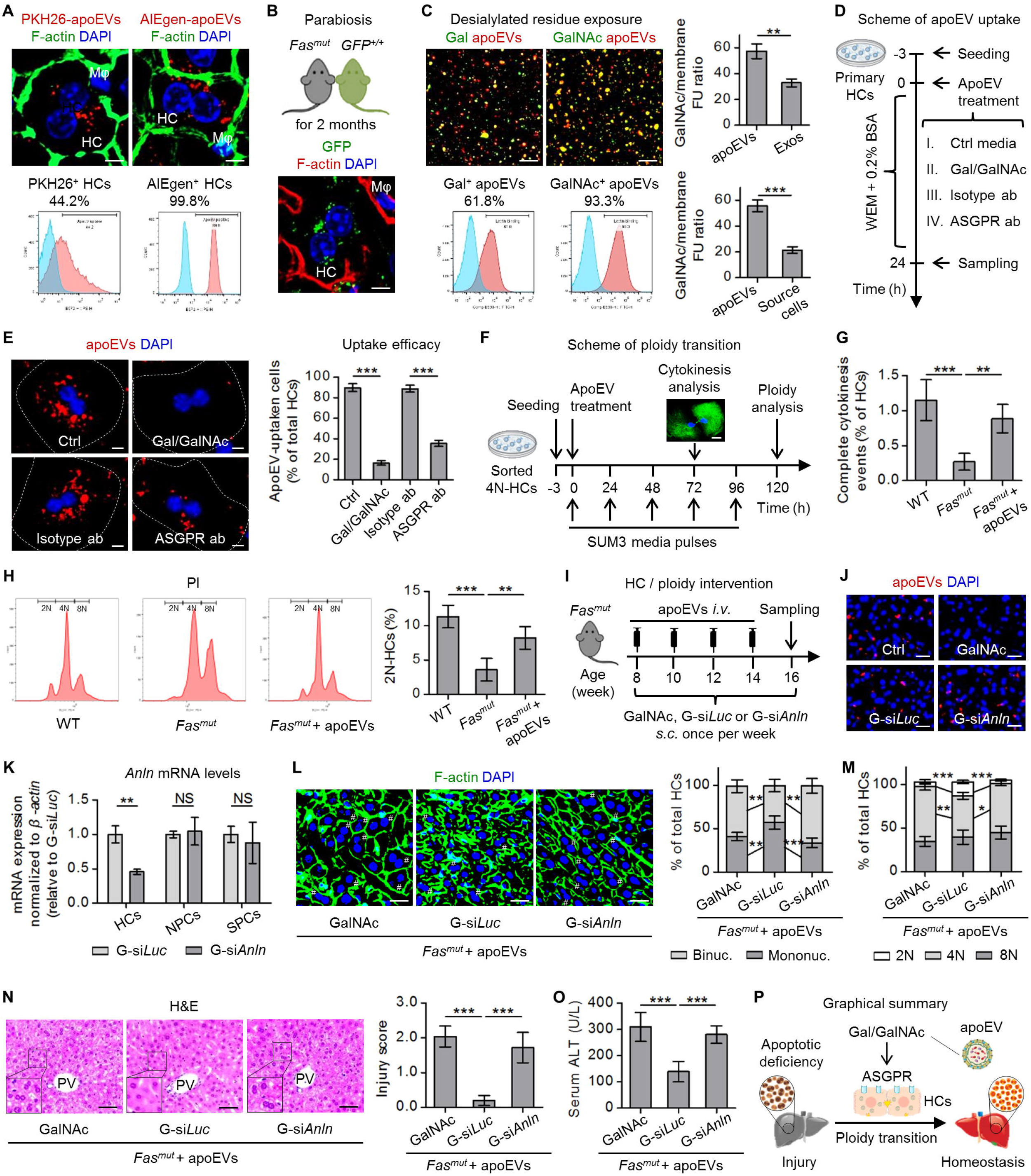
Specialized internalization of apoEVs by hepatocytes (HCs) promotes ploidy transition. **(A)** Tracing of PKH26-labeled (for membrane) or aggregation-induced emission luminogen (AIEgen, for mitochondria)-labeled apoEVs (red) in liver, with phalloidin staining F-actin (green) to show HC cell borders. DAPI was used for counterstaining DNA (blue). ApoEVs were intravenously injected and liver samples were collected 24 h after the infusion. Representative confocal images show binucleated HCs and the neighbored macrophages (Mφs). Flow cytometric analyses were performed on freshly isolated primary HCs to detect the fluorescent signals and determine percentages of apoEV-positive HCs. Scale bars = 10 μm. **(B)** Tracing of green fluorescent protein (GFP)-labeled particles (green) in liver of *Fas* mutant (*Fas^mut^*) mice, with phalloidin staining F-actin (red) to show borders of HC cells. DAPI was used for counterstaining DNA (blue). *Fas^mut^* mice and *GFP^+/+^* transgenic (*GFP’ “*) mice were connected to generate parabiosis model for 2 months. A representative confocal image shows a binucleated HC and a neighboring Mφ. Scale bar = 10 μm. **(C)** Representative fluorescent images showed apoEV surface exposure of desialylated residues including galactose (Gal) and *N*-acetylgalactosamine (GalNAc). ApoEVs (labeled by CellMask™ Deep Red, red) were stained with FITC-conjugated lectins (green) specific for Gal or GalNAc binding and aggregation. Flow cytometric analysis was performed to determine percentages of apoEV staining. ApoEV exposure of GalNAc was compared to exosomes (Exos) or to apoEV source mesenchymal stem cells (MSCs) using fluorescent unit (FU, analyzed by a fluorescent reader) over membrane labeling (by CellMask™ Deep Red) FU ratio. Scale bars = 25 μm. *N* = 3 per group. **(D)** Schematic diagram demonstrates the study design of apoEV uptake by primary cultured HCs. WEM, William’s E Medium; BSA, bovine serum albumin; ASGPR, asialoglycoprotein receptor; ab, antibody. ApoEVs were treated at a protein concentration of 10 μg/ml. Gal and GalNAc were added together at 200 mM each, and antibodies were added at 5 μg/ml. **(E)** Representative fluorescent images show PKH26-labeled apoEVs (red) uptaken by binucleated HCs (DAPI for DNA, blue). White dashed lines depict cell borders of cultured HCs. Flow cytometric analysis was performed to detect the fluorescent signals and determine the percentage of apoEV-positive HCs. Scale bars = 10 μm. *N* = 5 per group. **(F)** Schematic diagram demonstrates the study design of HC ploidy transition. ApoEV protein concentration was 10 μg/ml. Representative image shows complete cytokinesis, which is identified by the midbody structure formed by microtubules (MTs, α-tubulin immunostaining, green; counterstained with DAPI, blue). Scale bar = 20 μm. **(G)** Imaging analysis of complete cytokinesis events. WT, wild type. *N* = 4 per group. **(H)** After propidium iodide (PI) staining, ploidy distribution of HCs (diploid, 2N; tetraploid, 4N; octoploid, 8N) was analyzed by flow cytometry. *N* = 4 per group. **(I)** Schematic diagram demonstrating the study design of *in vivo* HC uptake or HC ploidy intervention upon apoEV infusion. ApoEVs were infused intravenously (*i.v.*) at a protein concentration of 5 μg/g. GalNAc was injected subcutaneously (*s.c.*) at 500 mg/kg once per week to inhibit apoEV uptaken by HCs. GalNAc-conjugated small interferon RNA (siRNA) against the *Anillin* gene (*G-siAnln*) for inhibiting HC cytokinesis or its luciferase control (*G-siLuc*) was injected at 4 μg/g. **(J)** Tracing of PKH26-labeled apoEVs (red) in the liver. DAPI was used for DNA counterstaining (blue). Scale bars = 25 μm. **(K)** Quantitative real-time polymerase chain reaction (qRT-PCR) analysis of *Anln* mRNA expression in freshly isolated HCs, liver non-parenchymal cells (NPCs) and splenocytes (SPCs) with *ß-actin* as internal control. *N* = 3 per group. **(L)** Representative liver fluorescent images show HCs with different numbers of nuclei (blue, DAPI for DNA) and their cell borders (green, phalloidin for F-actin). # indicates binucleated HCs. Percentages of binucleated and mononucleated HCs were quantified. Scale bars = 25 μm. *N* = 5 per group. **(M)** After PI staining, ploidy distribution of HCs was analyzed by flow cytometry. *N* = 3 per group. **(N)** Hematoxylin and eosin (H&E) staining of liver tissues in periportal vein (PV) areas. Hepatic injury scores were assessed based on pathological parameters. Scale bars = 50 μm. *N* = 5 per group. **(O)** Serum alanine aminotransferase (ALT) levels were determined. *N* = 5 per group. **(P)** Graphical summary demonstrates that apoEV uptake by HCs promotes ploidy transition for liver homeostasis. Data represent mean ± standard deviation. *, *P* < 0.05; **, *P* < 0.01; ***, *P* < 0.0001; NS, not significant, *P* > 0.05. Statistical analyses were performed using Student’s *t* test for two-group analysis or one-way analysis of variance followed by the Newman-Keuls post hoc tests for multiple group comparisons.

We continued to dissect the reasons for and functional results of apoEVs being selectively taken up by HCs. It has been documented that HCs specifically bind and internalize galactose (Gal)- or *N*-acetylgalactosamine (GalNAc)-terminating glycoproteins through the featured asialoglycoprotein receptor (ASGPR) (Ashwell and Harford, 1982), and that desialylated glycans characterized by Gal and GalNAc residues are exposed on the surface of apoptotic cells (Dini et al., 1992). Thus, we investigated whether apoEVs use the sugar recognition system for HC internalization. Using fluorescent lectin conjugates specifically binding the desialylated residues, we revealed that both Gal and GalNAc were exposed on the surface of apoEVs; particularly, GalNAc marked over 90% of apoEVs (Figure 2C). Furthermore, GalNAc abundance in apoEVs was higher than in Exos and in source MSCs when normalized by total membrane labeling fluorescent units, suggesting an apoEV-specific profile (Figure 2C). To evaluate whether the Gal/GalNAc-ASGPR system is indeed pivotal to apoEV uptake by HCs, we treated primary cultured HCs with apoEVs under different conditions with competing exogenous Gal/GalNAc or the ASGPR-specific antibody (Dini et al., 1992; McVicker et al., 2002) (Figure 2D). Data demonstrated that both the added monosaccharide mixture and the ASGPR antibody effectively inhibited HC uptake of apoEVs (Figure 2E). Furthermore, we noticed in this simple short-term culture process that mononucleated diploid HCs became binucleated and polyploidized, and we managed to track the subsequent ploidy transition of tetraploid hepatocytes in particular using a mitogenic stimulation system after sorting (Duncan et al., 2010) (Figure 2F). We discovered that 4N-HCs from *Fas^mut^* mice rarely underwent complete cytokinesis marked by the microtubule “midbody” structure (Figure 2G) and failed to produce 2N daughter cells (Figure 2H). Intriguingly, apoEV treatment empowered the *Fas^mut^* 4N-HCs to bypass cytokinesis failure and induced an emergence of 2N populations, indicating ploidy reversal (Figure 2G, H). Collectively, these data suggest that specialized uptake of apoEVs by HCs promotes ploidy transition.

The above results prompted us to investigate whether apoEV-promoted HC ploidy transition is of functional significance to the liver. By sorting and comparing properties of 2N- and 4N-HCs (Figure S4A), we discovered that 4N-HCs possessed lower oxidative metabolic activity but accumulated more reactive oxygen species (ROS) than 2N-HCs, irrespective of the derivation from apoEV deficiency or replenishment conditions (Figure S4B, C). The oxidative stress in 4N-HCs was possibly due to a general decline of expression of antioxidant genes (Figure S4D), potentially underlying the injury-proneness of binucleated HCs in the liver. Notably, apoEV treatment maintained the oxidative phosphorylation (OXPHOS) rate and restrained the oxidative stress of unsorted HCs (Figure S4E, F), which might be attributed to ploidy reversal effects of apoEVs.

To confirm whether apoEVs indeed regulate liver function by targeting HCs to promote ploidy transition, we designed *in vivo* experimental protocols using high-dose intermittent injection of GalNAc at 500 mg/kg to inhibit apoEV uptake by HCs, and low-dose intermittent administration of GalNAc-conjugated small interfering RNAs (siRNAs) for *Anillin* (G-si*Anln*) at 4 mg/kg to specifically inhibit HC cytokinesis (Figure 2I). As previously established, GalNAc conjugation allows targeted delivery of siRNAs to HCs with limited off-target effects, and Anillin is an actin-binding protein required for cytokinesis (Zhang et al., 2018a; Zhang et al., 2018b). We confirmed that only the high-dose GalNAc injection effectively suppressed liver engraftment of apoEVs (Figure 2J), and that G-si*Anln* administration selectively down-regulated *Anln* mRNA expression in HCs (Figure 2K). Furthermore, both GalNAc and G-si*Anln* administration inhibited apoEV restoration of HC ploidy homeostasis in *Fas^mut^* mice, leaving lower percentages of mononucleated and diploid HCs in the liver compared to the control siRNA-injected (G-si*Luc*) *Fas^mut^* mice with apoEV infusion (Figure 2L, M). Importantly, hematoxylin and eosin (H&E) staining of liver tissues and serum ALT analysis demonstrated that either blocking HC uptake of apoEVs or inhibiting HC cytokinesis diminished apoEV-mediated maintenance of liver integrity under apoptotic deficiency (Figure 2N, O). These results suggested that internalization of apoEVs by HCs promotes ploidy transition for liver homeostasis (Figure 2P).

We further examined the parabiosis model to investigate whether physiological apoEVs contribute to liver homeostasis maintenance (Figure S5A). Intermittent injection of GalNAc was also applied to inhibit HC uptake of apoEVs in *GFP^+/+^-paired Fas^mut^* mice (Figure S5A, B). As shown by H&E staining and serum ALT analysis, 2-month parabiosis with healthy *GFP^+/+^* mice helped the paired *Fas^mut^* mice to recover from liver injury, while GalNAc injection suppressed the effects (Figure S5C, D). Furthermore, parabiosis with *GFP^+/+^* mice reduced the percentages of binucleated HCs and promoted emergence of 2N-HCs in the liver of *Fas^mut^* mice, both of which were also counteracted by GalNAc injection (Figure S5E, F). These data indicate that circulatory endogenous apoEVs maintained liver ploidy and functional homeostasis through HC uptake.

### ApoEVs interact with recipient Golgi apparatus to form a chimeric organelle complex for HC cytokinesis

Next, we aimed to decipher how apoEVs facilitate ploidy transition of HCs. In the previous *in vivo* and *in vitro* tracing experiments, we noticed that internalized apoEVs gathered in the perinuclear region of HCs (Figure 2A and 2E). Intriguingly, it has been reported that following endocytosis, the Golgi apparatus receives cargo by a process called backward/retrograde transport (Johannes and Popoff, 2008), while the destination of EVs in recipient cells remains uncertain (Heusermann et al., 2016; Liu et al., 2020). We stained HC Golgi *in vivo* using an antibody against Golgin84 (also known as Golga5), an integral Golgi membrane protein. We discovered that almost all internalized apoEVs in binucleated HCs contact with the Golgi membrane, which occurred rapidly at 24 h post-apoEV infusion or as a physiological event during parabiosis (Figure 3A). To further characterize apoEV interactions with the recipient Golgi, we transfected sorted 4N-HCs in culture with GFP-plasmids labeling *N*-acetylgalactosaminyltransferase 2 (GALNT2), a Golgi-localized enzyme. We found that apoEVs were indeed closely adjacent to and specifically contacted the perinuclear Golgi apparatus in interphase, during which apoEVs and recipient Golgi formed a chimeric three-dimensional structure by apoEVs “inserting” into Golgi “pits”, “bridging” Golgi tubules and circling around Golgi “ends” (Figure 3B). The Golgi apparatus is increasingly acknowledged as a microtubule organizing center (MTOC) which safeguards mitotic spindle formation and cell division (Wei et al., 2015). In addition, microtubules are regulated by post-translational modifications, including acetylation at the lysine 40 (K40) residues of α-tubulin, which indicates stabilization of microtubules for functional execution (Montagnac et al., 2013). We further revealed that perinuclear interactions between apoEVs and Golgi in binucleated HCs were also associated with microtubule organization, particularly formation of acetylated microtubules, which also enriched in the perinuclear region and co-localized with the apoEV-Golgi chimeric structure in the interphase (Figure 3C). Moreover, the intimate spatial and structural correlations among apoEVs, Golgi and acetylated microtubules were also detected in HC cytokinesis with the midbody actually formed by the acetylated microtubules (Figure 3C), suggesting functional contributions to the cytokinesis process.

**Figure 3.**
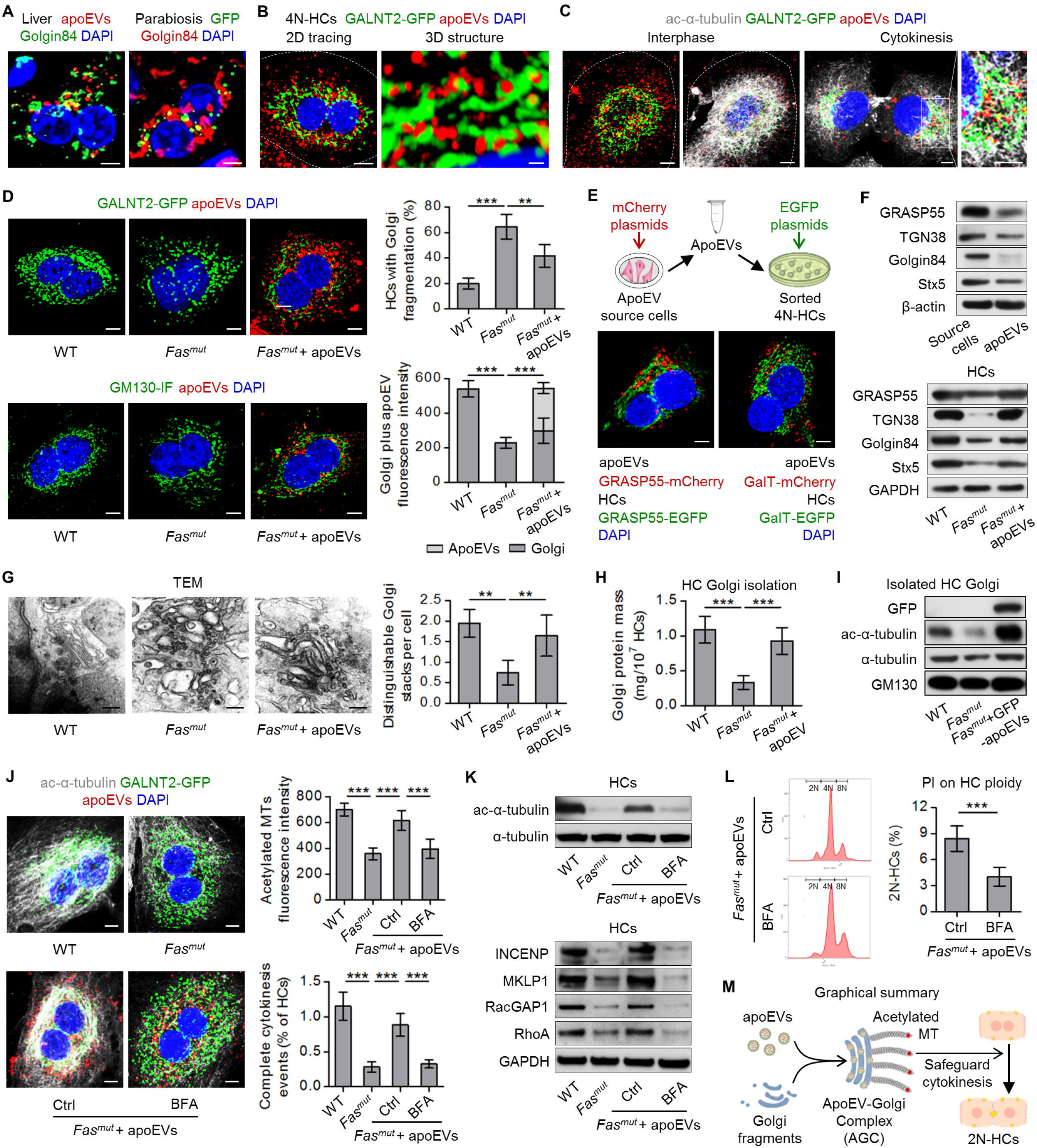
ApoEVs interact with recipient Golgi apparatus to form a chimeric organelle complex. **(A)** Left: Tracing of aggregation-induced emission luminogen (AIEgen)-labeled apoEVs (red) in liver, with Golgin84 immunostaining for Golgi apparatus (green) and counterstaining with DAPI (blue). ApoEVs were intravenously injected and liver samples were collected at 24 h after the infusion. Right: Tracing of green fluorescent protein (GFP)-labeled particles (green) in the liver of *Fas* mutant (*Fas^mut^*) mice, with Golgin84 immunostaining for Golgi apparatus (red) and counterstaining with DAPI (blue). *Fas^mut^* mice and *GFP* transgenic (*GFP^+/+^*) mice were connected as a parabiosis model for 2 months. Representative confocal images show binucleated HCs. Scale bars = 5 μm. **(B)** Tracing of PKH26-labeled apoEVs (red) in sorted tetraploid HCs (4N-HCs), with Golgi apparatus being demonstrated by a plasmid of *N*-acetylgalactosaminyltransferase 2 (GALNT2)-GFP (green) and counterstaining with DAPI (blue). Representative 2D confocal images show binucleated HCs and 3D reconstructed images show apoEV contacting Golgi apparatus. Scale bars = 10 μm (2D) and 500 nm (3D). **(C)** Representative fluorescent images show cultured 4N-HCs in interphase and in cytokinesis with GALNT2-GFP-labeled Golgi (green), PKH26-labeled apoEVs (red), DAPI counterstain (blue), and ac-α-tubulin immunostaining for acetylated microtubules (MTs, white). Scale bars = 10 μm. **(D)** Representative fluorescent images show cultured 4N-HCs with GALNT2-GFP-labeled Golgi (green, up) or 130 kDa Golgi matrix protein (GM130) immunofluorescent (IF) staining (green, down), PKH26-labeled apoEVs (red), and DAPI counterstain (blue). Imaging analysis was performed to quantify HC percentages with Golgi fragmentation and Golgi with apoEV fluorescence intensity. WT, wild type. Scale bars = 5 μm. *N* = 4 per group. **(E)** Schematic diagram demonstrates the study design for investigating apoEV-Golgi assimilation. Representative fluorescent images show cultured 4N-HCs with their own Golgi proteins (green) and apoEV-transferred Golgi proteins (red) with DAPI counterstain (blue). GRASP55, Golgi reassembly-stacking protein 55; GalT, galactosyltransferase. Scale bars = 5 μm. **(F)** Western blot analysis shows multiple Golgi protein expression in apoEVs compared to their source MSCs, and in cultured HCs. TGN38, *trans-Golgi* network protein 38; Stx5, syntaxin 5; GAPDH, glyceraldehyde 3-phosphate dehydrogenase. **(G)** Representative transmission electron microscopy (TEM) images show Golgi ultrastructure in sorted 4N-HCs. Distinguishable Golgi stacks per cell were quantified. Scale bars = 200 nm. *N* = 4 per group. **(H)** Golgi apparatus was isolated from cultured HCs and Golgi protein mass was determined using the BCA method. *N* = 4 per group. **(I)** Western blot analysis showed GFP expression, MT and acetylated MT markers in isolated Golgi. ApoEVs were derived from *GFF^+/+^* MSCs. GM130 was used as an internal control. **(J)** Representative fluorescent images show cultured 4N-HCs with ac-α-tubulin immunostaining for acetylated MTs (white), GALNT2-GFP-labeled Golgi (green), PKH26-labeled apoEVs (red), and DAPI counterstain (blue). Imaging analysis was performed to quantify acetylated MTs fluorescence intensity and complete cytokinesis events. Ctrl, control; BFA, Brefeldin A, added at 1 nM. Scale bars = 5 μm. *N* = 4 per group. **(K)** Western blot analysis show MT and expression of multiple mitosis proteins in cultured HCs. INCENP, inner centromere protein; MKLP1, mitotic kinesin-like protein 1; RacGAP1, Rac GTPase-activating protein 1; RhoA, Ras homolog family member A. **(L)** After propidium iodide (PI) staining, ploidy distribution in HCs (diploid, 2N; tetraploid, 4N; octoploid, 8N) was analyzed by flow cytometry. *N* = 4 per group. **(M)** Graphical summary demonstrating apoEV assembly with Golgi to safeguard HC cytokinesis. Data represent mean ± standard deviation. **, *P* < 0.01; ***, *P* < 0.0001. Statistical analyses were performed by Student’s *t* test for two-group analysis or one-way analysis of variance followed by the Newman-Keuls post hoc tests for multiple group comparisons.

We were particularly interested in the chimeric structure formed by apoEVs and recipient Golgi, and further evaluated whether this interaction had functional implications for Golgi maintenance. By comparing the GALNT2-labeled Golgi morphology in sorted 4N-HCs from WT and *Fas^mut^* mice, we unexpectedly discovered that whereas the Golgi in WT 4N-HCs exhibited a relatively complete ribbon structure, the Golgi in *Fas^mut^* 4N-HCs was fragmented and dispersed (Figure 3D). Notably, when tracing labeled apoEVs in *Fas^mut^* 4N-HCs, the apoEVs enriched in the perinuclear region, where a complete Golgi was ought to be, while maintaining contact with the Golgi fragments or as Golgi “replenishments” (Figure 3D). As the GALNT2 represents a *medial/trans-Golgi* marker, we also performed immunofluorescent (IF) staining of 130 kDa Golgi matrix protein (GM130), a *cis*-Golgi marker. Data demonstrated that the GM130 fluorescence was also dispersed in *Fas^mut^* 4N-HCs, whereas in *Fas^mut^* 4N-HCs after apoEV treatment, the GM130-labeled structure was tighter, with internalized apoEVs embedding among the Golgi stacks (Figure 3D). ApoEV contacts with the Golgi and protection of Golgi integrity were also found in *Casp3^-/-^* 4N-HCs (Figure S3H). These results, especially the apoEV-Golgi structural assimilation, inspired us to hypothesize that integration of apoEVs with the recipient Golgi may form a vesicle-organelle complex.

To test this hypothesis, we transfected apoEV source cells (*i.e.* MSCs) and sorted 4N-HCs with mCherry- or EGFP-plasmids for enforced expression of the same targets, Golgi reassembly-stacking protein 55 (GRASP55, responsible for stacking of Golgi cisternae located in *medial-*Golgi) and GalT (galactosyltransferase, an enzyme on *trans-*Golgi membranes) (Figure 3E). After collecting apoEVs from the transfected MSCs and treating HCs with the apoEVs, we detected donor GRASP55/GalT integrated with their recipient counterparts (Figure 3E). Indeed, apoEVs carried a spectrum of Golgi proteins with varied abundance, including the Golgi stacking protein GRASP55, the *trans-*Golgi network protein 38 (TGN38), Golgin84 for tethering of vesicles, and Syntaxin 5 (Stx5) for docking and fusion of vesicles (Figure 3F). Correspondingly, *Fas^mut^* HCs derived from apoptosis-deficient conditions showed varying degree of decline in the expression of multiple Golgi proteins, while apoEV treatment in culture replenished the insufficient Golgi proteins (Figure 3F). Rescue of Golgi integrity was further confirmed under electron microscopy, which revealed that the Golgi in *Fas^mut^* 4N-HCs was mostly dispersed into vesicle forms, whereas distinguishable Golgi stacks were observed after apoEV treatment (Figure 3G). Moreover, when quantified after isolation from cultured HCs, Golgi in the apoEV-deficient *Fas^mut^* mice lost over 50% of the protein mass, which was recovered by apoEV replenishment (Figure 3H). ApoEV-mediated Golgi recovery was verified in *Casp3^-/-^* HCs as well (Figure S3I). Also, after using apoEVs derived from *GFP^+/+^* MSCs, the GFP signal was detected in the isolated HC Golgi (Figure 3I). Taken together, these data suggest that integration of apoEVs with recipient Golgi apparatus safeguarded the structural integrity of the Golgi organelle.

We then examined the functional results downstream of apoEV-induced Golgi recovery. First, we confirmed that either α-tubulin or its acetylated form (ac-α-tubulin) was detected in the isolated HC Golgi (Figure 3I), indicating the Golgi was structurally conjugated to microtubules. Notably, *Fas^mut^* HC Golgi showed a decreased ac-α-tubulin level compared to WT HC Golgi, while apoEV treatment promoted the Golgi-conjugated microtubule acetylation (Figure 3I). These protein expression changes on isolated HC Golgi were consistent with general cellular changes of the 4N-HCs, in which apoEV treatment rescued the diminished perinuclear acetylated microtubules together with the assembled Golgi against *Fas^mut^* apoptotic deficiency (Figure 3J). To further demonstrate the significance of Golgi recovery for mediating effects of apoEVs, we applied Brefeldin A (BFA), a chemical that is commonly used to disrupt the Golgi organization. We revealed that BFA treatment induced Golgi disassembly despite the perinuclear apoEVs, and it also diminished apoEV-promoted acetylated microtubules and inhibited apoEV-rescued cytokinesis in *Fas^mut^* 4N-HCs (Figure 3J). Western blot analyses of microtubule acetylation levels in HCs confirmed these results, showing apoEV-upregulated ac-α-tubulin expression in *Fas^mut^* HCs being suppressed by BFA (Figure 3K). Moreover, apoEV application triggered upregulation of a series of downstream cascade proteins for cytokinesis (Barr and Gruneberg, 2007) in *Fas^mut^* HCs (Figure 3K), such as the inner centromere protein (INCENP, a part of the chromosomal passenger complex regulating cytokinesis), the mitotic kinesin-like protein 1 (MKLP1, also known as KIF23, a motor protein forming the centralspindlin complex with Racgap1), Rac GTPase-activating protein 1 (RacGAP1, a microtubule-dependent signal for Rho regulation), and finally Ras homolog family member A (RhoA, the master regulator of contractile ring formation for abscission). Positive effects of apoEVs on the cytokinesis proteins were also diminished by BFA treatments (Figure 3K), resulting in suppressed function of apoEVs to promote 2N-HC emergence during ploidy transition of *Fas^mut^* 4N-HCs (Figure 3L). Collectively, the above data indicated that apoEVs assembled with recipient Golgi apparatus to form a chimeric organelle structure, which we hereby name as the ApoEV-Golgi Complex (AGC), for driving microtubule acetylation and safeguarding the cytokinesis process of HCs (Figure 3M).

### ApoEVs use the SNARE mechanism to assemble with Golgi and maintain liver homeostasis

Next, we investigated the molecules responsible for AGC assembly. The soluble *N*-ethylmaleimide-sensitive factor attachment protein receptor (SNARE) mechanism is recognized as mediating the final step in vesicle transport, *i.e.* docking and fusion of vesicles with the target membrane compartment, which is dependent on the interactions between specific molecules on the vesicles (the v-SNAREs) and on the target membranes (the t-SNAREs) (Pfeffer, 1996). Currently, there are three known v-SNAREs that mediate endosomal interactions with the Golgi in the retrograde route of transport: vesicle-associated membrane protein 3 (VAMP3, also known as cellubrevin), VAMP4 and Golgi SNARE of 15 kDa (GS15) (Johannes and Popoff, 2008). With this knowledge, we examined whether apoEVs, as examples of extracellular vesicles, may adopt the intracellular vesicle/endosomal mechanisms to interact with the Golgi apparatus. IF staining and flow cytometric analyses demonstrated that high percentages of apoEVs expressed VAMP3 and VAMP4 on their surfaces (84.0% positive for VAMP3, 66.7% positive for VAMP4), but virtually none expressed GS15 (less than 10% positive) (Figure 4A). Furthermore, VAMP3 protein expression in apoEVs was even higher than in the source MSCs (Figure 4B). To evaluate whether VAMP3 and/or VAMP4 indeed mediate the AGC assembly, we transfected source MSCs with siRNAs for *Vamp3* and/or *Vamp4* and collected the *Vamp*-knockdown apoEVs for functional experiments (Figure 4C). Data demonstrated that in *Fas^mut^* 4N-HCs, knockdown of *Vamp3* alone was enough to prevent apoEVs from docking with the recipient Golgi without affecting the uptake, and the internalized *Vamp3*-knockdown apoEVs scattered in the cytoplasm away from the Golgi (Figure 4D). Knockdown of *Vamp4* had limited effects on the fate of apoEVs in recipient cells (Figure 4D). We further revealed that *Vamp3*-knockdown apoEVs did not rescue the fragmented Golgi, nor did they restore the diminished acetylated microtubules in *Fas^mut^* 4N-HCs (Figure 4E). To directly confirm that apoEVs adopted VAMP3 to assemble with the Golgi, we used the isolated Golgi system and detected hVAMP3 when treating HCs with hApoEVs, indicating integration of apoEV-VAMP3 into the recipient Golgi (Figure 4F). These data suggest that VAMP3 was the key factor in apoEV-mediated AGC assembly and functional effects of apoEVs.

**Figure 4.**
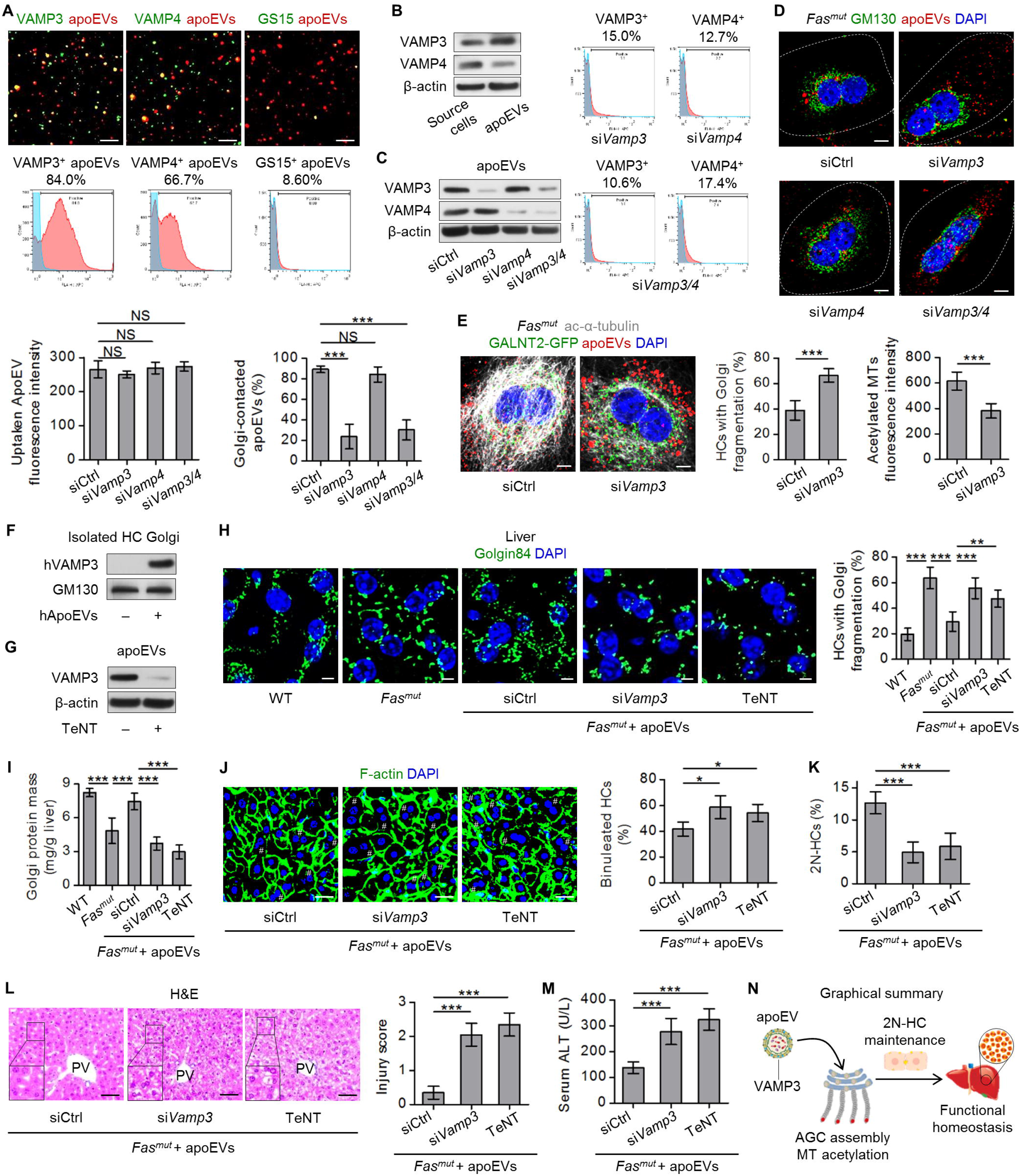
ApoEVs use vesicle-associated membrane protein 3 (VAMP3) to assemble with Golgi. **(A)** Representative immunofluorescent images show apoEV (labeled with CellMask™ Deep Red, red) surface expression of vesicle-localized-soluble *N*-ethylmaleimide-sensitive factor attachment protein receptors (v-SNAREs, green) VAMP3, V VAMP4 and Golgi SNARE of 15 kDa (GS15). Flow cytometric analysis was performed to determine percentages of positively stained apoEVs. Scale bars = 25 μm. **(B)** Western blot analysis show expression of VAMPs in apoEVs compared to their source MSCs. **(C)** Western blot analysis show expression of VAMPs in apoEVs. Source MSCs were transfected with small interferon RNAs (siRNAs) as a negative control (siCtrl) for VAMP3 (si*Vamp3*), VAMP4 (si*Vamp4*), and both VAMP3 and VAMP4 (si*Vamp3/4*). Flow cytometric analysis was performed to determine percentages of positively stained apoEVs after knockdown of VAMPs. **(D)** Representative images show cultured *Fas* mutant (*Fas^mut^*) tetraploid hepatocytes (4N-HCs) with 130 kDa Golgi matrix protein (GM130) immunofluorescent staining (green), PKH26-labeled apoEVs (red), and DAPI counterstain (blue). White dashed lines depict cell borders of cultured HCs. Imaging analysis was performed to quantify apoEV uptake and percentages of Golgi-contacted apoEVs. Scale bars = 10 μm. *N* = 4 per group. **(E)** Representative fluorescent images show cultured *Fas^mut^* 4N-HCs with ac-α-tubulin immunostaining (white), *N*-acetylgalactosaminyltransferase 2 (GALNT2)-GFP-labeled Golgi (green), PKH26-labeled apoEVs (red), and DAPI counterstain (blue). Imaging analysis was performed to quantify HC percentages with Golgi fragmentation and acetylated microtubules (MTs) fluorescence intensity. Scale bars = 5 μm. *N* = 4 per group. **(F)** Western blot analysis showed human VAMP3 (hVAMP3) expression in isolated HC Golgi. ApoEVs were derived from human MSCs (hApoEVs). GM130 was used as an internal control. **(G)** Western blot analysis showed VAMP3 expression in apoEVs without or with treatment by 250 ng/ml Tetanus toxin (TeNT). **(H)** Representative Golgin84 immunofluorescent staining (green) images of liver Golgi counterstained with DAPI (blue). WT, wild type. Imaging analyses were performed to quantify HC percentages with Golgi fragmentation. Scale bars = 5 μm. *N* = 4 per group. **(I)** Golgi apparatus was isolated from the liver and Golgi protein mass was determined using the BCA method. *N* = 4 per group. **(J)** Representative liver fluorescent images showed HCs with different nuclei (blue, DAPI for DNA) and their cell borders (green, phalloidin for F-actin). # indicates binucleated HCs. Scale bars = 25 μm. *N* = 4 per group. **(K)** After PI staining, percentages of binucleated HCs were quantified. Diploid HCs (2N-HCs) were analyzed using flow cytometry. *N* = 4 per group. **(L)** Hematoxylin and eosin (H&E) staining of liver tissues in periportal vein (PV) areas. Hepatic injury scores were examined based on pathological parameters. Scale bars = 50 μm. *N* = 4 per group. **(M)** Serum alanine aminotransferase (ALT) levels were determined. *N* = 4 per group. **(N)** Graphical summary illustrating that apoEVs use VAMP3 to assemble with Golgi for HC and liver regulation. Data represent mean ± standard deviation. *, *P* < 0.05; **, *P* < 0.01; ***, *P* < 0.0001; NS, not significant, *P* > 0.05. Statistical analyses were performed by Student’s *t* test for two-group analysis or one-way analysis of variance followed by the Newman-Keuls post hoc tests for multiple group comparisons.

It has been reported that certain VAMPs, including VAMP3, on synaptic vesicles can be cleaved by Tetanus toxin (TeNT) (McMahon et al., 1993). We verified that VAMP3 on apoEVs was also sensitive to TeNT treatment (Figure 4G), which provided a useful tool for evaluating apoEV-VAMP3 function *in vivo*. Importantly, as shown by Golgin84 IF staining, we discovered that *in vivo* binucleated HCs in *Fas^mut^* mice were fragmented, consistent with the dispersed Golgi of *Fas^mut^* 4N-HCs in culture (Figure 4H). Furthermore, total Golgi protein mass in *Fas^mut^* mouse liver was decreased compared to WT mouse liver, corresponding to the decreased Golgi protein mass in cultured *Fas^mut^* HCs (Figure 4I). Notably, chronic intermittent infusion of apoEVs partially rescued the defective HC Golgi together with the declined liver Golgi in *Fas^mut^* mice (Figure 4H, I). These defects were also rectified by long-term parabiosis with *GFP^+/+^* mice, based on GalNAc-mediated physiological apoEV internalization (Figure S5G, H). Expectedly, infusion of *Vamp3*-knockdown apoEVs or TeNT-pretreated VAMP3-cleaved apoEVs failed to restore Golgi integrity in *Fas^mut^* mice (Figure 4H, I), confirming VAMP3 as the critical mediator of apoEV effects. The significance of VAMP3 to apoEV function was further verified by *Vamp3* knockdown or TeNT pretreatment abolishing the promotion of HC ploidy reversal and protection against liver injury in *Fas^mut^* mice by infused apoEVs (Figure 4J-M). Taken together, these results indicate that apoEVs used the SNARE mechanism to assemble with Golgi and activate downstream functional cascades to maintain liver homeostasis (Figure 4N).

### ApoEV-mediated Golgi recovery and ploidy reversal contribute to liver regeneration

Next, we investigated the implications of apoEV regulation of the liver. The liver is well known for its remarkable regenerative capabilities, which initiates a coordinated series of mechanisms to regain its original volume following injury or resections (Forbes and Newsome, 2016). Interestingly, it has been documented that HC proliferation and/or hypertrophy contribute to liver regeneration after PHx, while rapid division of binucleated HCs without mitosis particularly underlies liver regeneration after 70% PHx, a large-scale surgery after which the liver can recover (Hori et al., 2011; Mitchell and Willenbring, 2008; Miyaoka et al., 2012). Moreover, a recent study has demonstrated that the polyploid state restricts HC proliferation and liver regeneration in mice (Wilkinson et al., 2019b), although controversies exist (Matsumoto et al., 2020; Wilkinson et al., 2019a; Zhang et al., 2018a). We applied the commonly used 70% PHx model to examine the participation of apoEVs in the liver regeneration process (Figure 5A). At 72 h after 70% PHx, when one round of HC cell cycle is typically finished (Sakamoto et al., 1999), we confirmed regeneration of the remnant liver tissues to over 60% of the original liver mass, thus guaranteeing the survival of the mice (Figure 5B). Notably, compared to sham-treated mice that only underwent laparotomy, PHx mice demonstrated an increase in Annexin V-labeled hepatic apoEVs (Figure 5C), which might be related to a wave of cellular apoptosis accompanying mitosis of HCs (Sakamoto et al., 1999), indicating apoEVs being actively involved in the liver regeneration process. Furthermore, we discovered a slight increase of Golgin84-stained area in the regenerating liver (Figure 5D), together with a reduced percentage of binucleated HCs (Figure 5E), suggesting responses of Golgi being associated with apoEV changes and the ploidy transition of HCs in liver regeneration.

**Figure 5.**
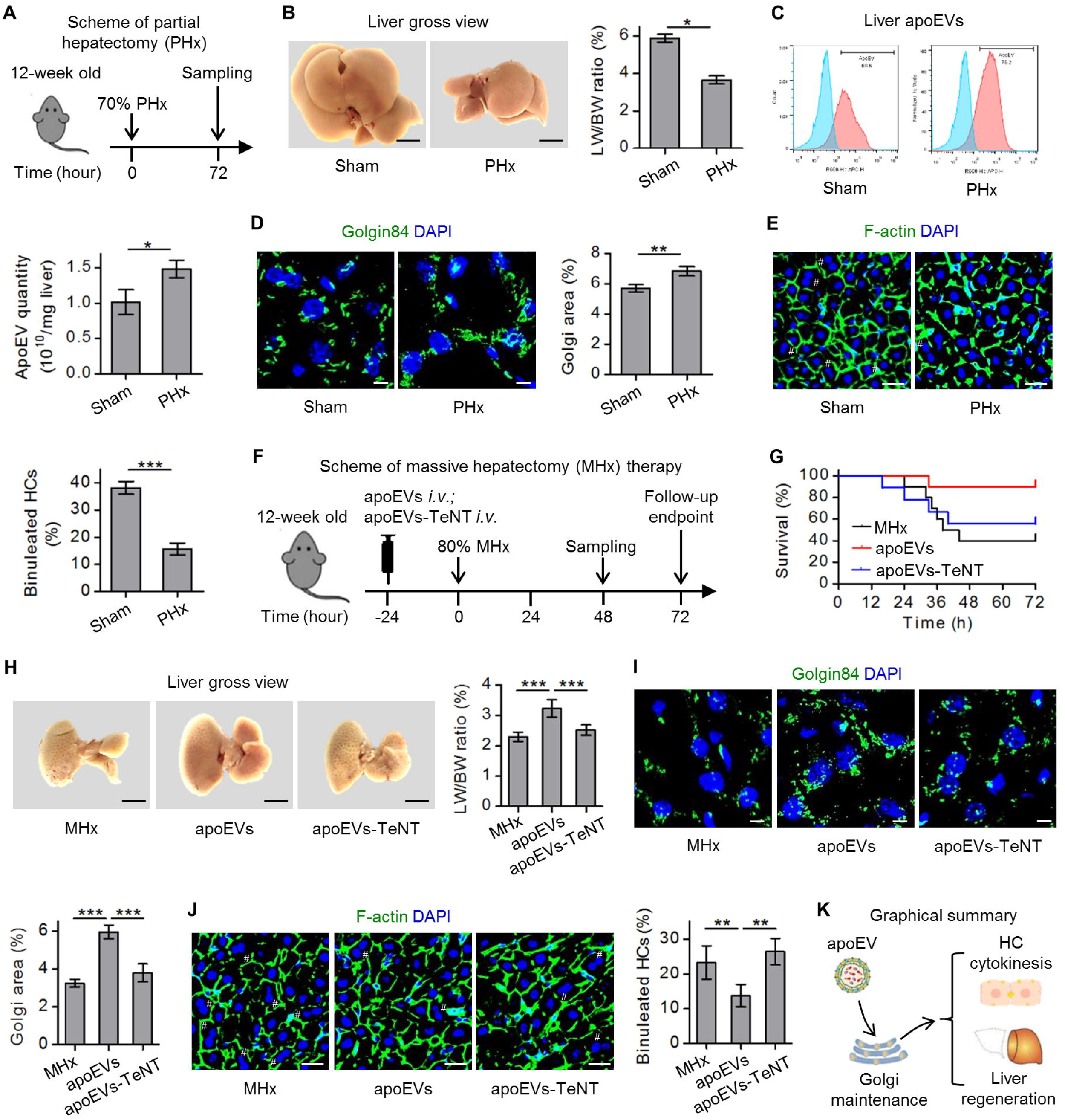
ApoEV-mediated Golgi recovery and ploidy reversal contribute to liver regeneration. **(A)** Schematic diagram demonstrates the study design of 70% partial hepatectomy (PHx). **(B)** Gross view images of liver and quantification of liver weight (LW) to body weight (BW) ratio after recovery for 72 h. Scale bars = 5 mm. *N* = 5 per group. **(C)** Flow cytometric analysis showed Annexin V^+^ liver tissue extracellular vesicles (apoEVs) and corresponding quantification. *N* = 3 per group. **(D)** Representative Golgin84 immunofluorescent staining (green) images of liver Golgi counterstained with DAPI (blue). Imaging analysis was performed to quantify Golgi area percentages. Scale bars = 5 μm. *N* = 4 per group. **(E)** Representative liver fluorescent images show hepatocytes (HCs) with different numbers of nuclei (blue, DAPI for DNA) and their cell borders (green, phalloidin for F-actin). # indicates binucleated HCs. Percentages of binucleated HCs were accordingly quantified. Scale bars = 25 μm. *N* = 4 per group. **(F)** Schematic diagram demonstrates the study design of 80% massive hepatectomy (MHx) therapy. ApoEVs at a protein concentration of 5 μg/g were infused intravenously (*i.v.*) alone or after administration of 250 ng/ml Tetanus toxin (TeNT). **(G)** Kaplan-Meier survival curve of MHx mice. *N* = 10 per group. **(H)** Gross view images of the liver and quantification of LW/BW ratio after recovery for 48 h. Scale bars = 5 mm. *N* = 5 per group. **(I)** Representative Golgin84 immunofluorescent staining (green) images of liver Golgi counterstained with DAPI (blue). Imaging analysis was performed to quantify Golgi area percentages. Scale bars = 5 μm. *N* = 4 per group. **(J)** Representative liver fluorescent images show HCs with different numbers of nuclei (blue, DAPI for DNA) and their cell borders (green, phalloidin for F-actin). # indicates binucleated HCs. Percentages of binucleated HCs were quantified. Scale bars = 25 μm. *N* = 4 per group. **(K)** Graphical summary demonstrates apoEVs promote Golgi recovery and ploidy reversal for liver regeneration. Data represent mean ± standard deviation. *, *P* < 0.05; **, *P* < 0.01; ***, *P* < 0.0001. Statistical analyses were performed by Student’s *t* test for two-group analysis or one-way analysis of variance followed by the Newman-Keuls post hoc tests for multiple group comparisons.

To reveal the functional significance of apoEVs to liver regeneration, we established an 80% massive hepatectomy (MHx) mouse model (Figure 5F). Without intervention, mice may not survive this injury and typically die within 72 h after surgery (Hori et al., 2011). Importantly, infusion of apoEVs at 24 h prior to surgery substantially improved the survival rate of MHx mice (Figure 5G) and promoted recovery of liver mass at 48 h after surgery (Figure 5H). Infusion of TeNT-pretreated VAMP3-cleaved apoEVs failed to promote liver regeneration and improve survival of MHx mice (Figure 5G, H). As expected, infusion of apoEVs significantly upregulated the Golgi abundance in liver after MHx, and the effects were diminished by TeNT pretreatments (Figure 5I). Moreover, apoEV infusion accelerated the division of binucleated HCs for rapid liver regeneration dependent on apoEV-VAMP3 expression (Figure 5J). Collectively, these data provide proof-of-concept evidence that apoEV-mediated Golgi recovery and ploidy reversal contribute to liver regeneration (Figure 5K).

### ApoEVs protect against lethal liver failure through Golgi and ploidy restorations

The above findings encouraged us to further unravel the therapeutic potential of apoEVs in counteracting liver diseases, by applying a clinically relevant model of ALF induced by lethal acetaminophen (APAP) challenge (Barbier-Torres et al., 2017; Stravitz and Lee, 2019). Due to overdoses of paracetamol and other medications for pain and fever control, APAP toxicity represents the principal cause of ALF in developed countries, estimated in the United States at about 30,000 patients hospitalized per year and accounting for 45.7% of all ALF cases (Blieden et al., 2014; Stravitz and Lee, 2019). However, how liver regeneration becomes overwhelmed or impaired in this circumstance, thus failing to restore the severely damaged liver, is not fully understood (Forbes and Newsome, 2016). In previous experiments, we noticed that declined hepatic apoEVs in *Fas^mut^* mice and *Casp3^-/-^* mice led to the onset of liver injury, which did not heal until apoEVs were replenished. We further observed in the lethally APAP-damaged liver, which again especially demonstrated severely injured binucleated HCs (Figure 6A-C), that hepatic apoEVs were reduced (Figure 6D). Furthermore, we also found impaired liver Golgi after APAP challenge (Figure 6E), whereas the regenerative division of binucleated cells was not initiated (Figure 6F). These findings indicate that apoEV exhaustion with Golgi and ploidy transition impairments underlie diminished liver regeneration in ALF.

**Figure 6.**
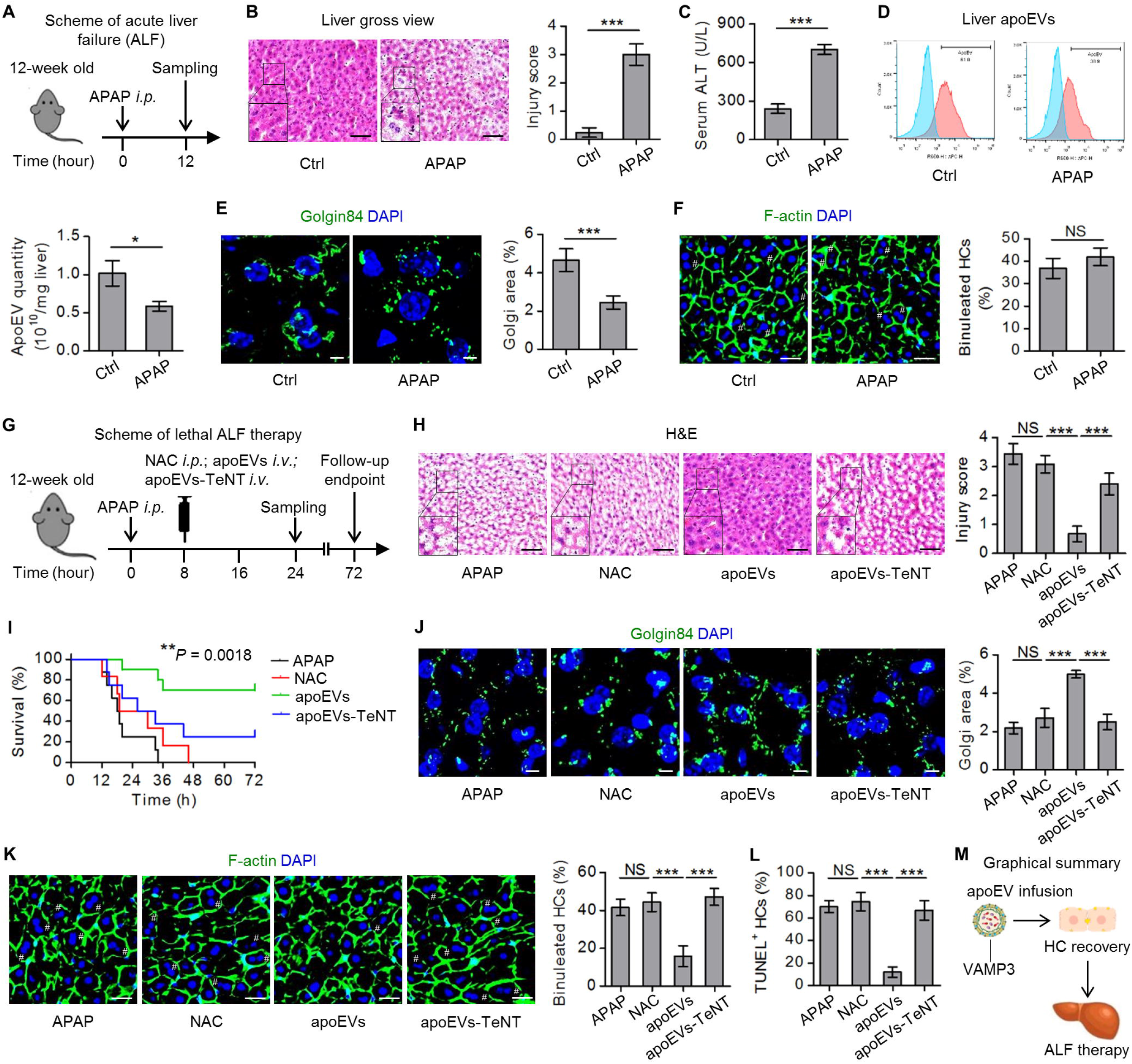
ApoEVs protect against acute acetaminophen (APAP)-induced liver failure through Golgi and ploidy restorations. **(A)** Schematic diagram demonstrates the study design of APAP-induced acute liver failure (ALF). APAP was injected intraperitoneally (*i.p.*) at 1 g/kg. **(B)** Hematoxylin and eosin (H&E) staining of liver tissues. Hepatic injury scores were examined based on pathological parameters. Ctrl, control. Scale bars = 100 μm. *N* = 5 per group. **(C)** Serum alanine aminotransferase (ALT) levels were determined. *N* = 5 per group. **(D)** Flow cytometric analysis of Annexin V^+^ liver tissue extracellular vesicles (apoEVs) and corresponding quantification. *N* = 3 per group. **(E)** Representative Golgin84 immunofluorescent staining (green) images of liver Golgi counterstained with DAPI (blue). Imaging analysis was performed to quantify Golgi area percentages. Scale bars = 5 μm. *N* = 4 per group. **(F)** Representative liver fluorescent images show hepatocytes (HCs) with different numbers of nuclei (blue, DAPI for DNA) and their cell borders (green, phalloidin for F-actin). # indicates binucleated HCs. Percentages of binucleated HCs were quantified. Scale bars = 25 μm. *N* = 5 per group. **(G)** Schematic diagram demonstrates the study design of ALF therapy. APAP was injected *i.p.* at 1 g/kg. ApoEVs at protein quantification of 5 μg/g were infused intravenously (*i.v.*) with or without 250 ng/ml Tetanus toxin (TeNT). The antioxidant *N*-acetyl L-cysteine (NAC) was injected *i.p.* at 1 g/kg. **(H)** H&E staining of liver tissues. Hepatic injury scores were examined based on pathological parameters. Scale bars = 100 μm. *N* = 5 per group. **(I)** Kaplan-Meier survival curve of ALF mice. *N* = 10 per group. **(J)** Representative Golgin84 immunofluorescent staining (green) images of liver Golgi counterstained with DAPI (blue). Imaging analysis was performed to quantify Golgi area percentages. Scale bars = 5 μm. *N* = 4 per group. **(K)** Representative liver fluorescent images show HCs with different numbers of nuclei (blue, DAPI for DNA) and their cell borders (green, phalloidin for F-actin). # indicates binucleated HCs. Percentages of binucleated HCs were quantified. Scale bars = 25 μm. *N* = 4 per group. **(L)** Percentages of apoptotic HCs were quantified by terminal deoxynucleotidyl transferase-mediated dUTP nick-end labeling (TUNEL) staining. *N* = 4 per group. **(M)** Graphical summary demonstrates apoEV infusion promotes HC recovery for ALF therapy. Data represent mean ± standard deviation. *, *P* < 0.05; **, *P* < 0.01; ***, *P* < 0.0001; NS, not significant, *P* > 0.05. Statistical analyses were performed by Student’s *t* test for two-group analysis, one-way analysis of variance followed by the Newman-Keuls post hoc tests for multiple group comparisons, or the Log-rank test for survival curve comparisons.

We therefore explored whether apoEV infusion could serve as an effective therapeutic for ALF (Figure 6G). Currently, the antioxidant *N*-acetyl L-cysteine (NAC) is the sole available and standard therapy to treat APAP overdose patients, but it is only effective as an antidote within a short time window of less than 8-12 h after APAP ingestion (Barbier-Torres et al., 2017). In light of this information, we applied apoEV infusion at 8 h post-APAP challenge and compared it to NAC supplied at the same time point (Figure 6G). Data demonstrated that while NAC administration at this time failed to rescue the severe liver damage induced by APAP, apoEVs provoked a surprisingly complete liver protection in surviving mice at 24 h after APAP injection (Figure 6H). Also, the effects of apoEVs were abolished by TeNT pretreatment for VAMP3 cleavage (Figure 6H). Further evaluation of mouse survival revealed that apoEV infusion, but not NAC injection, dramatically lowered the death rates, with effects diminished by TeNT pretreatment (Figure 6I). The ability of apoEVs to ameliorate APAP-induced ALF were attributed to the rescue of liver Golgi and further emergence of mononucleated HCs in liver, together with declined HC apoptosis, which were suppressed after apoEV-VAMP3 being cleaved by TeNT (Figure 6J-L). Collectively, these results suggest that apoEV infusion promotes HC recovery for ALF therapy (Figure 6M).

We took a final step to examine whether the functional molecules we discovered on the surface of STS-induced MSC-derived apoEVs, *i.e.* Gal/GalNAc and VAMP3, were general profiles of apoEVs as functional markers for liver regulation. Based on flow cytometric analyses established above, we tested the exposure of aopEVs to PtdSer (Annexin V-binding), Gal/GalNAc and VAMP3 in the following circumstances: human MSCs (hMSCs) treated with ABT-737, a B-cell lymphoma-2 (Bcl-2) homology domain 3 (BH3)-mimetic that directly triggers permeabilization of the mitochondrial outer membrane to induce the intrinsic pathway of apoptosis (Medina et al., 2020) (Figure S6A); mouse activated T cells with Fas-crosslinking (anti-Fas treatment) to induce the extrinsic pathway of apoptosis, which represents a physiological source of apoEVs (Figure S6B); and STS-treated NIH/3T3 cells, which are a mouse fibroblast cell line representing differentiated cells, another physiological source of apoEVs (Figure S6C). Our data demonstrated that apoEVs collected from the different conditions uniformly exposed PtdSer and were characterized by surface Gal, GalNAc and VAMP3 (Figure S6A-C), suggesting a general apoptotic vesicular signature (AVS) marking a functional “envelope” of apoEVs for liver regulation (Figure S6D).

## Discussion

Apoptosis is recognized as an integral physiological cell death process which occurs daily with a high prevalence and generates a huge number of apoEVs (Arandjelovic and Ravichandran, 2015; Atkin-Smith et al., 2015; Caruso and Poon, 2018; Fuchs and Steller, 2011). However, whether apoEVs are necessary for maintaining organ homeostasis has remained unclear. In this study, we aimed to investigate the biological properties and the related physiological function of apoEVs in organ maintenance. We characterized apoEVs by their specific AVS and possession of liver engraftment properties. We found that apoEVs were recognized and internalized by HCs in the liver through the Gal/GalNAc-ASGPR system, in which they assembled with recipient Golgi apparatus to form a chimeric AGC structure based on the SNARE mechanism. AGC formation in binucleated HCs then activated downstream functional cascades including microtubule acetylation and cytokinesis, which were indispensable for ploidy transition of HCs to safeguard the diploid populations. Ultimately, apoEVs were of functional significance for maintaining liver homeostasis, promoting liver regeneration and mediating therapy against ALF. These findings collectively suggest a previously unrecognized role and mechanism of apoEVs in regulating liver function, shedding light on the potential of developing apoEV therapy for liver diseases.

Apoptosis represents the dominant modality of cell death in the normal homeostasis of multicellular organisms, and is indispensable not only for necessary cell elimination but also for regulation of fundamental life processes such as immune reactions, metabolism and regenerative proliferation (Fuchs et al., 2013; Fuchs and Steller, 2011; Han and Ravichandran, 2011; Morioka et al., 2019). Effects of apoptosis are exerted through either a “passive” mode by efferocytosis of apoptotic “corpses”, during which macrophages exert their own function (Morioka et al., 2018), or a more “active” form that involves controlled release of nucleotides, proteins and metabolites for regulation of phagocytosis and compensatory proliferation (Chekeni et al., 2010; Chera et al., 2009; Medina et al., 2020; Wickman et al., 2013). Among the paracrine signals, apoEVs possess unique properties as subcellular membrane-delimited particles, which encapsulate heterogeneous factors for distant and combinational messaging (Atkin-Smith et al., 2015; Dieude et al., 2015; Liu et al., 2018). We have previously reported that circulating apoEVs maintain stem cell function in bone and ameliorate osteopenia *via* transferring concerted signaling modulators (Liu et al., 2018). Others have documented that apoEVs deliver multiple mitogenic or alarm cues to trigger molecular and functional cascades for epithelial turnover and inflammation (Berda-Haddad et al., 2011; Brock et al., 2019; Wickman et al., 2013). In the present study, we established a regulatory link between hepatic/systemic apoEVs and liver homeostasis, mediated by specific apoptotic vesicular markers for destined engraftment and functional internalization. ApoEV contributions to liver homeostasis further underlie the effects of organismal apoptotic regulation. As far as we know, this is the first evidence for the requirement of apoEVs in organ homeostatic maintenance and reveals the previously unknown, specific mechanisms involved. The heterogeneity of apoEVs introduced by source cells, during the apoptotic process and among the potential subsets of apoEVs will be a critical issue to be addressed in next studies. Future studies will also need to elaborate upon the physiological source and identity of liver-regulating apoEVs, and trace their involvement in liver homeostasis and pathologies.

The liver is a vital organ that functions as the primary center for sustaining metabolism and disposing of toxins, and it is also known to clear systemic apoptotic cells through its resident macrophages (Shi et al., 1996). However, as apoptotic materials may overwhelm the phagocytic capacity of liver macrophages (Morioka et al., 2019), it is plausible that other hepatic cell populations contribute to digestion of the apoptotic “meal”. As the major cell type in the liver responsible for its function, HCs account for about 80% of liver weight and about 70% of all liver cells; HCs also show the ability to engulf apoptotic cells (Dini et al., 1992; McVicker et al., 2002). In this study, notably, we reveal that HCs actually represent the main target of apoEVs in the liver, particularly for those free of xenobiotic membrane labeling. Furthermore, internalization of apoEVs by HCs induces functional results such as promoting cytokinesis. One possible mechanism enlightened by this study is that apoEVs control Golgi structure formation and function, as intact Golgi promotes microtubule polymerization and organization through the containing of γ-tubulin, the major nucleator of microtubules (Rios et al., 2004), and α-tubulin acetyltransferase 1 (αTAT1), the acetyltransferase specific to the K40 residue of α-tubulin (Nakakura et al., 2016), and improved Golgi-derived microtubule organization facilitates cell division and cytokinesis (Wei et al., 2015). These findings provide a new perspective on cell death, indicating that the contents of vesicles produced during apoptosis are naturally required for functional purposes, rather than just being efferocytosed. Intriguingly, other EV populations (*e.g.* Exos) also engraft in the liver and are involved in liver regulation (Wiklander et al., 2015; Ying et al., 2017), but the underlying reasons why the liver needs EVs are still elusive. Here, we for the first time uncovered that apoEVs indispensably participate in the unique ploidy transition process of HCs, which is characteristically seen in hepatic postnatal development, aging and pathologies (Fortier et al., 2017; Wang et al., 2017). It is notable that non-apoptotic EVs may emerge to maintain a constant liver EV pool upon loss of apoEVs, as indicated in this study, whereas the endogenous source of the non-apoptotic EVs and their functional roles remain unrecognized. Howbeit, our findings contribute to explaining the high demand of the liver for apoEVs at the functional level, indicating the apoptotic materials reciprocally potentiate dynamic behaviors of living cells. These data consistent with recent studies by us and others suggesting that diploid HCs play a key role in maintaining liver homeostasis and regeneration (Gentric et al., 2015; Wang et al., 2014; Wilkinson et al., 2019b), although they are relatively low in abundance and their functional roles are still in controversial (Kreutz et al., 2017; Zhang et al., 2018a). Moreover, whether apoEVs exert differential effects on diploid and polyploid HCs, and whether apoEVs realize ploidy reversal of mononucleated polyploid HCs, remain to be investigated. Also to be examined is the responses of aneuploid HCs to apoEVs, which might promote genetic diversity among HCs in the liver and facilitate stress-induced liver adaptation (Duncan et al., 2012; Duncan et al., 2010). Our results pave an avenue for further extensive examinations of apoEV regulation of general HC and liver function, including but not limited to the age-related polyploidization process, the glucose/lipid metabolism and the protein/enzyme synthesis.

The eukaryotes possess an ancient and elaborate endomembrane system which is responsible for the biosynthesis and transport of proteins and lipids, as well as for the formation of major intracellular organelles (Field and Dacks, 2009). The endomembrane system is dynamically maintained by endosomal vesicular trafficking in a bi-directional mode: anterograde transport ends with exocytosis and retrograde transport starts with endocytosis, while the Golgi serves as the crucial interface between these two processes (Johannes and Popoff, 2008; Lee et al., 2004). Interestingly, EVs, as products of endosomal or plasma membrane origin, also have close relationships with the endomembrane system for trafficking after internalization (Heusermann et al., 2016; Joshi et al., 2020). In a previous study, we have reported that apoEVs after engulfment regulate recipient lysosomal biogenesis by activating a master transcription factor (Liu et al., 2020). However, it remains unknown whether EVs or apoEVs directly integrate into recipient organelles with functional significance. In this study, we discovered that apoEVs assemble with recipient Golgi and are required for Golgi integrity, which adds new dimensions to the current knowledge in the following respects. First, homeostasis of an organelle or at least a part of the endomembrane system demands extrinsic EV inputs. Second, EVs can be integrally reused as necessary structural units for functional execution. Finally, the Golgi apparatus might actually be a chimeric organelle (or the AGC structure) in certain cells with physiological apoEV internalization. One mechanism revealed in this study is that apoEVs express v-SNAREs and apoEVs use VAMP3 to assemble with Golgi. According to current models, as far as we know, SNARE-mediated membrane contacts may end up with different outcomes, such as a full-collapse fusion or just a kiss-and-run mode of flicker fusion (Smith et al., 2008). Although VAMP3 functionally expresses on apoEVs for finding Golgi, how apoEVs work with Golgi will still be interesting to further examine. Whether other EVs also assemble with recipient organelles and whether apoEVs fuse with endosomes or other membranous organelles remain to be investigated. It is intriguing findings that Golgi integrity is required for liver homeostasis and that Golgi participates in liver regeneration, whereas loss of Golgi contributes to ALF progression.

Although the regulatory mechanisms of Golgi assembly and disassembly in organ health and sickness are not fully understood, the involvements of Golgi in stress responses and disease pathogenesis emerge to be noticed (Machamer, 2015; Sbodio et al., 2018). In this regard, apoEV infusion may serve as an effective approach to potentiate healthy Golgi for incoming functional requirements *(e.g.* in MHx) or rescue defective Golgi (*e.g.* in ALF). Moreover, as the Golgi apparatus performs multiple important functions such as protein sorting and modification (Lee et al., 2004), it would be significant to decipher the potential role of the Golgi as a cargo- or signaling-processing platform for internalized EVs, which might be linked to the undissected mechanisms apoEVs in promoting microtubule acetylation *via* the Golgi in this study.

To cope with a constant influx of noxious stimuli, the liver has developed a remarkable and tightly controlled regenerative capacity in response to parenchymal loss, which nevertheless can be impaired in severe or chronic injury settings (Forbes and Newsome, 2016). Thus, liver disease is a primary cause of mortality and morbidity that is also rising globally (Williams et al., 2017). Currently, for end-stage liver disease or failure, the only clinical approach is liver transplantation, but availability is limited by several major challenges including a donor shortage (Stravitz and Lee, 2019). Therefore, there remains an unmet need to rescue patients from lethal liver injuries. In this study, on the basis of previous findings on apoptotic regulation of tissue regeneration (Brock et al., 2019; Li et al., 2010), we unraveled that infusion of allogeneic apoEVs significantly promotes liver regeneration after PHx and can serve as an effective therapeutic for ALF. ApoEV therapy holds great promise for translational applications, especially when the current antidote fails to work in late-stage ALF. Based on the results of this study, we assume that by apoEVs promoting ploidy transition, the ROS toxicity-prone 4N-HCs are rapidly replaced by 2N-HCs for liver protection, while whether this is a form of action of detoxification or regeneration will be tricky to explain. It is notable that with translational advantages, therapeutic apoEVs can be easily obtained at a larger amount than Exos and without ultracentrifuge from cultured MSCs, which possess immense potential for regenerative and modulatory transplantation and have been used extensively in clinical trials for treating a variety of diseases (Galipeau and Sensebe, 2018). Furthermore, given the potential hepatotrophic property of apoEVs, engineered chimeric apoEVs can be constructed for targeted delivery of therapeutic molecules or drugs, as we previously established (Dou et al., 2020), which might prove useful for treating complicated chronic diseases of the liver. Nevertheless, the application of apoEVs in treating liver diseases should also be cautious particularly considering that the polyploid state may play a tumor-suppressive role in the liver (Zhang et al., 2018a; Zhang et al., 2018b). Although that apoEVs are kind of safeguarding ploidy homeostasis of the liver without promoting diploid HCs over the physiological status, how apoEVs might balance the effects and risks remain elusive. Taken together, our findings open a window for future translational research on apoEVs to counteract diseases in the liver context.

## Acknowledgements

This work was supported by grants from the Guangdong Financial Fund for High-Caliber Hospital Construction (174-2018-XMZC-0001-03-0125, D-07 to S.S., D-11 to X.K.), the Pearl River Talent Recruitment Program (2019ZT08Y485), the National Science and Technology Major Project of the Ministry of Science and Technology of China (2018ZX10302207-001-002), the Sun Yat-sen University Young Teacher Key Cultivation Project (18ykzd05 to X.K.), the National Natural Science Foundation of China (32000974 to B.S.), the Postdoctoral Innovative Talents Support Program of China (BX20190380 to B.S.), and the General Program of China Postdoctoral Science Foundation (2019M663986 to B.S.). We thank Alnylam’s Bioinformatics, Oligo Synthesis, and RNAi Lead Discovery team for designing, generating, and screening for the Anln GalNAc siRNAs. We thank Prof. Zhiying He in Tongji University for discussion on liver ploidy analysis. We thank Dr. Zijian Jiang in The Fourth Military Medical University for technical support on hepatectomy surgeries. We thank Dr. Allyson J Merrell and Prof. Ben Z Stanger in University of Pennsylvania for technical support on culture of primary HCs.

## Author contributions

B.S. contributed to the study design and conduct, and the manuscript drafting. R.W. contributed to data acquisition, analysis and interpretation. C.C. and X.K. contributed to data analysis and interpretation. D.W. contributed to hepatectomy experiments. H.C.T. contributed to GalNAc-based hepatocyte ploidy manipulation. Y.W. contributed to the experimental design on Golgi apparatus. Y.L., O.J. and X.C. contributed to labeling of apoEVs by radionuclides.

H.L., R.T.K.K. and B.Z.T. contributed to labeling of apoEVs by the AIE luminogen. H.Y. contributed to design and interpretation of liver experiments. M.W. contributed to design and interpretation of ploidy experiments. L.X. and X.Y. contributed to apoEV identification and marker analysis. Y.F. contributed to apoptosis verification. X.Z., J.T. and L.M. contributed to data interpretation. L.L. contributed to mouse breeding and profiling. Y.J. and S.S. contributed to the project conception, experimental design and supervision. All authors contributed to the manuscript revision and approved the final version of the manuscript.

## METHODS

### LEAD CONTACT AND MATERIALS AVAILABILITY

Further information and requests for resources and reagents should be directed to and will be fulfilled by the Lead Contact, Songtao Shi.

### METHOD DETAILS

#### Animals

The following mouse strains were obtained from the Jackson Laboratory: *Fas^mut^* (B6.MRL-*Fas^lpr^*/J, JAX# 000482), *Casp3^-/-^* (B6N.129S1-*Casp3^tm1Flv^*/J, JAX# 006233, provided as heterozygotes), *GFP^+/+^* (C57BL/6-Tg(CAG-EGFP)1Osb/J, JAX# 003291), and C57BL/6J (JAX# 000664) as the WT (Liu et al., 2018). Male mice at ages of 8 or 12 weeks were used. All mice were housed in pathogen-free conditions, maintained on a standard 12-h light-dark cycle, and received food and water *at libitum.* Genotyping was performed by polymerase chain reaction (PCR) using tail samples from mice and primer sequences provided by the Jackson Laboratory. All animal experiments were performed in compliance with the relevant laws, institutional guidelines and ARRIVE guidelines.

#### Reagents

All antibodies, chemicals, cytokines and plasmids used in this study are listed in the Key Resources Table (Table S1).

#### Isolation and culture of MSCs

Mouse MSCs were isolated from hind limb bone marrow and cultured, as we previously described (Liu et al., 2018). Briefly, a single suspension of bone marrow cells was seeded at a density of 1.5×10^7^ cells per 10 cm culture dish at 37°C in 5% CO_2_. Non-adherent cells were removed after 24 h and attached cells were maintained for 16 days in alpha-Minimum Essential Medium (α-MEM) (12571-048, Invitrogen, USA) supplemented with 20% fetal bovine serum (FBS) (17L624, Sigma-Aldrich, USA), 2 mM L-glutamine (35050-061, Invitrogen, USA), 55 μM 2-mercaptoethanol (21985-023, Invitrogen, USA), 100 U/ml penicillin (15140-122, Invitrogen, USA) and 100 μg/ml streptomycin (15140-122, Invitrogen, USA). Colony-forming attached cells were passed twice by trypsin (12605-010, Invitrogen, USA) for further experimental use.

Human MSCs were isolated from full-term umbilical cords and cultured, as we previously described (Sun et al., 2010). Briefly, fresh umbilical cords were obtained from informed, healthy mothers in local maternity hospitals of China after normal deliveries and were processed under sterile conditions. The cords were rinsed twice in phosphate buffered saline (PBS) in penicillin and streptomycin, and outer membrane and vessels were removed. The washed cords were then cut into 1 mm^2^ pieces and incubated in low-glucose Dulbecco’s modified Eagle’s medium (DMEM) (11885-084, Invitrogen, USA) containing 10% FBS (S12450, Atlanta Biologicals, USA), 2 mM L-glutamine (35050-061, Invitrogen, USA), 100 U/ml penicillin (15140-122, Invitrogen, USA) and 100 μg/ml streptomycin (15140-122, Invitrogen, USA) at 37°C in a humidified atmosphere of 5% CO_2_. Colony-forming attached cells were passed by trypsin (12605-010, Invitrogen, USA) for further experimental use.

#### Isolation and culture of T lymphocytes

Mouse T cells were derived from spleen and cultured, as we previously described (Sui et al., 2017). Briefly, with ACK Lysing Buffer (10-548E, Lonza Bioscience, Switzerland) was used to remove red blood cells. T cells were isolated and stimulated for 48 h with 3 μg/ml plate-bound anti-mouse CD3 antibody (100340, BioLegend, USA) and 2 μg/ml soluble anti-mouse CD28 antibody (102116, BioLegend, USA) in RPMI 1640 medium (21870-076, Invitrogen, USA) supplemented with 10% FBS (17L624, Sigma-Aldrich, USA), 2 mM L-glutamine (35050-061, Invitrogen, USA), 100 U/mL penicillin (15140-122, Invitrogen, USA), and 100 g/ml streptomycin (15140-122, Invitrogen, USA) at 37°C in a humidified atmosphere of 5% CO_2_.

#### Culture of the cell line

NIH/3T3 cell line was purchased from ATCC and was maintained in low-glucose DMEM (11885-084, Invitrogen, USA) supplemented with 10% FBS (17L624, Sigma-Aldrich, USA), 2 mM L-glutamine (35050-061, Invitrogen, USA), 100 U/ml penicillin (15140-122, Invitrogen, USA) and 100 μg/ml streptomycin (15140-122, Invitrogen, USA) at 37°C in a humidified atmosphere of 5% CO_2_.

#### Induction and examination of apoptosis

Apoptosis was induced by multiple methods in this study. To induce cell apoptosis *via* both caspase-dependent and caspase-independent pathways, as we previously established (Liu et al., 2018), STS (ALX-380-014, Enzo Life Sciences, USA) was added to serum-free medium at 500 nM for 16 h. For oxidative stress-induced apoptosis, as we previously established with minor modifications (Zheng et al., 2018), mitochondrial photodynamic activation was performed on AIEgen-stained cells in serum-free conditions with ultralow-power light irradiation at 0.7 mW/cm^2^ for 6 h. To initiate apoptosis *via* the intrinsic pathway, as reported (Medina et al., 2020), the BH3-mimetic ABT-737 (T2099, TargetMol, USA) which directly induces permeabilization of the mitochondrial outer membrane was applied at 5 μM for 48 h. To initiate apoptosis *via* the extrinsic pathway, as previously reported (Medina et al., 2020), T cells were treated with the CD95/Fas antibody (152803, BioLegend, USA) at 5 μg/ml for Fas-crosslinking during the 48 h activation. Apoptosis was identified by TUNEL (G3250, Promega, USA) or Annexin V (BMS500FI-300, Invitrogen, USA) / 7AAD (559925, BD Pharmingen, USA) staining.

#### Isolation, treatment and infusion of apoEVs

ApoEVs were isolated from medium of apoptotic cells using a sequential centrifugation method followed by filtering, as we previously established with modifications (Liu et al., 2018). Briefly, after sequential centrifugation at 800 g for 10 min at 4°C and 2,000 g for 10 min at 4°C followed by filtering through 5 μm filters, apoptotic cell lysates and debris were removed. Then, apoEVs were pelleted by centrifugation at 16,000 g for 30 min at 4°C and were diluted in PBS. Quantification of apoEVs was performed using the nanoparticle tracking analysis (NTA) for particle numbers and the BCA method (23225, Thermo Scientific, USA) for protein amounts. For treatment of HCs, apoEVs were added to the culture medium at a protein concentration of 10 μg/ml, with the dosage being determined according to preliminary tests. For intravenous infusion, apoEVs in PBS were injected *via* the mouse caudal vein at a protein concentration of 5 μg/g. For TeNT treatment of apoEVs to cleave VAMP3, TeNT (582243, Merck Millipore, Germany) was added to the apoEV solution at 250 ng/ml and incubated at 37°C for 30 min. ApoEVs were then centrifuged at 16,000 g for 30 min and washed twice with PBS before resuspension.

#### Labeling of apoEVs

ApoEVs were labeled using multiple methods in this study. ^64^Cu radionucleotide labeling of apoEVs was performed based on modifying our previous method of labeling of liposomes (Sun et al., 2017). Briefly, apoEVs were pelleted and resuspended in metal-free Na_2_HPO_4_ solution (pH = 7.5), into which NOTA-NHS ester (C100, CheMatech, France), a metal chelator, was added at 150 nM and incubated at 4°C overnight. Purification of the conjugated apoEVs was performed by centrifugation and washed with the Na_2_HPO_4_ solution. ^64^Cu labeling was performed by adding 20-30 mCi ^64^CuCl_2_ and the mixture was shaken at 37°C for 30 min followed by purification. The labeling efficiency was confirmed by instant thin-layer chromatography (ITLC) plates with citric acid (0.1 M, pH = 5) as an eluent. For labeling of apoEVs by membrane dyes, either PKH26 (PKH26PCL, Sigma-Aldrich, USA) or CellMask™ Deep Red (C10046, Invitrogen, USA) was used according to the manufacturers’ instructions. For labeling of apoEVs by the AIEgen, DCPy was synthesized according to our established protocol (Zheng et al., 2020). DCPy was then added to serum-free cell culture medium at a final concentration of 5 μM and incubated for 30 min followed by washes. Cells were then induced apoptosis by light irradiation, as stated above.

#### Isolation of Exosomes

Exosomes (Exos) were isolated from cultured un-apoptotic cells according to previous protocols with minor modifications (Liu et al., 2015). Briefly, cells were cultured in Exo-depleted medium (complete medium depleted of FBS-derived Exos by overnight centrifugation at 120,000 g) for 48 h. Exos from culture supernatants were isolated by differential centrifugation: 800 g for 10 min, 2,000 g for 10 min, 16,000 g for 30 min and 120,000 g for 120 min. Quantification of Exos was performed using the BCA method (23225, Thermo Scientific, USA) for protein amounts.

#### Isolation, sorting, culture and ploidy transition of primary HCs

Isolation of primary mouse HCs was performed according to previous protocols with minor modifications (Li et al., 2016). Briefly, mouse liver was perfused *via* catheterization of the portal vein using a 24G needle catheter (381212, BD, USA) and a mini-pump machine (13-876-2, Thermo Fisher, USA) under general anesthesia. The liver was perfused firstly with 10 ml Hank’s balanced salt solution (HBSS) (14175-079, Invitrogen, USA) to remove blood followed by 20 ml HBSS supplemented with 1 mM ethylene glycolbis(aminoethylether)-tetra-acetic acid (EGTA) (E3889, Sigma-Aldrich, USA) to remove the endogenous calcium. Then the liver was perfused with 20 ml HBSS supplemented with 5 mM CaCl_2_ (223506, Sigma-Aldrich, USA) and 40 μg/ml liberase TM (LIBTM-RO, Sigma-Aldrich, USA) for digestion. All the solutions were kept at 37°C in a water bath. After digestion, the liver was dissected and washed in ice-chilled HBSS, and cells were teased out into high-glucose DMEM (11965-084, Invitrogen, USA) supplemented with 10% FBS (17L624, Sigma-Aldrich, USA), 100 U/ml penicillin (15140-122, Invitrogen, USA) and 100 μg/ml streptomycin (15140-122, Invitrogen, USA). HCs were then prepared by centrifugation at 50 g for 5 min at 4°C, filtered through 70 μm nylon strainers, purified by a 49% Percoll solution (P4937, Sigma-Aldrich, USA), and resuspended in William’s E Medium (WEM) with GlutaMAX™ (32551-087, Invitrogen, USA) containing 10% FBS (17L624, Sigma-Aldrich, USA), 10 nM dexamethasone (D4902, Invitrogen, USA), 100 U/ml penicillin (15140-122, Invitrogen, USA) and 100 μg/ml streptomycin (15140-122, Invitrogen, USA). HCs were then seeded onto collagen-coated dishes (354450, Corning, USA) or coverslips (72295-14, Electron Microscopy Sciences, USA) and incubated at 37°C in a humidified atmosphere of 5% CO_2_ for 3 h for attachment.

For sorting of primary HCs, according to the published protocol (Duncan et al., 2010), isolated HCs before seeding were resuspended at 2×10^6^/ml for 30 min at 37°C in high-glucose DMEM (11965-084, Invitrogen, USA) supplemented with 10% FBS (17L624, Sigma-Aldrich, USA), 10 mM Hydroxyethyl piperazineethanesulfonic acid (HEPES) (H4034, Sigma-Aldrich, USA), 15 μg/ml Hoechst 33342 (H3570, Invitrogen, USA) and 5 μM reserpine (R0875, Sigma-Aldrich, USA). The HCs were then sorted for diploid and tetraploid HCs by flow cytometry and were collected in high-glucose DMEM (11965-084, Invitrogen, USA) supplemented with 50% FBS (17L624, Sigma-Aldrich, USA) and 10 mM HEPES (H4034, Sigma-Aldrich, USA). Sorted HCs were pelleted by centrifugation, resuspended and seeded in WEM with GlutaMAX™ (32551-087, Invitrogen, USA) containing 10% FBS (17L624, Sigma-Aldrich, USA), 10 nM dexamethasone (D4902, Invitrogen, USA), 100 U/ml penicillin (15140-122, Invitrogen, USA) and 100 μg/ml streptomycin (15140-122, Invitrogen, USA).

For apoEV uptake assays in HC culture, after attachment, HCs were washed twice with HBSS (14175-079, Invitrogen, USA) and medium was changed to WEM with GlutaMAX™ (32551-087, Invitrogen, USA) containing 0.2% bovine serum albumin (BSA) (700-100P, Gemini Bio, USA), 100 U/ml penicillin (15140-122, Invitrogen, USA) and 100 μg/ml streptomycin (15140-122, Invitrogen, USA) for following experiments (Li et al., 2016). ApoEVs were added at a protein concentration of 10 μg/ml for 24 h, during which 200 mM Gal (G5388, Sigma-Aldrich, USA) together with 200 mM GalNAc (HY-33212, MedChemExpress, China), or 5 μg/ml antibody for ASGPR (sc-166633, Santa Cruz Biotechnology, USA) or its isotype, were added to inhibit sugar recognition, as stated previously with minor modifications (McVicker et al., 2002).

Ploidy transition of cultured HCs was induced in a defined SUM3 medium (Duncan et al., 2010). SUM3 medium was prepared with 75% high-glucose DMEM (11965-084, Invitrogen, USA), 25% Waymouth’s medium (11220-035, Invitrogen, USA), 2 mM L-glutamine (35050-061, Invitrogen, USA), 100 U/ml penicillin (15140-122, Invitrogen, USA), 100 μg/ml streptomycin (15140-122, Invitrogen, USA), 10 mM HEPES (H4034, Sigma-Aldrich, USA), 50 ng/ml epidermal growth factor (EGF) (E4127, Sigma-Aldrich, USA), 1 μg/ml insulin (I0516, Sigma-Aldrich, USA), 30 nM sodium selenite (S5261, Sigma-Aldrich, USA), 10 μg/ml transferrin (T8158, Sigma-Aldrich, USA), 50 ng/ml somatotropin (869008-M, Sigma-Aldrich, USA) and 1 μM liothyronine (T_3_) (16028, Cayman Chemical, USA). After attachment, sorted 4N-HCs were washed twice with HBSS and medium was changed to the SUM3 medium, which was refreshed each day for 5 days. According to a protocol established through previous observations (Duncan et al., 2010; Fortier et al., 2017), cytokinesis was analyzed at 72 h by counting the microtubule midbody events, and ploidy status was analyzed at 120 h after collection by trypsin (12605-010, Invitrogen, USA). ApoEVs were added at a protein concentration of 10 μg/ml, and BFA (11861, Cayman Chemical, USA) was applied at 1 nM.

#### Isolation of liver NPCs, PBMNCs and BMMNCs

For isolation of liver NPCs, after liberase perfusion and cell collection, liver cells were centrifuged at 50 g for 5 min at 4°C to remove HCs, and the supernatant was centrifuged at 500 g for 5 min at 4°C to pellet NPCs. For PBMNC isolation, whole peripheral blood was extracted from the mouse retro-orbital venous plexus, and cells were isolated by centrifugation at 500 g for 5 min at 4°C followed by being treated with ACK Lysing Buffer (10-548E, Lonza Bioscience, Switzerland) to remove red blood cells. After washing with PBS, PBMNCs were collected by centrifugation at 500 g for 5 min at 4°C. For BMMNC isolation, whole bone marrow cells were treated with ACK Lysing Buffer (10-548E, Lonza Bioscience, Switzerland), washed with PBS, and collected by centrifugation at 500 g for 5 min at 4°C.

#### Isolation of liver tissue EVs and quantification

Liver tissue EV isolation was performed according to a previous protocol with modifications (Ishiguro et al., 2019). Briefly, after liberase perfusion and tissue collection, HCs were removed by centrifugation at 50 g for 5 min at 4°C, and the supernatant was centrifuged at 500 g for 5 min at 4°C to pellet NPCs. The supernatant of this step was further centrifuged at 2,000 g for 10 min at 4°C to remove tissue debris, filtered through 5 μm filters, and centrifuged at 16,000 g for 30 min at 4°C to pellet EVs. Quantifications of liver tissue EVs were performed using the NTA for particle numbers and apoEVs were detected by Annexin V-APC labeling (640920, BioLegend, USA).

#### Transfection of plasmids and siRNAs

Transfection of plasmids and siRNAs was performed according to our previous experience (Liu et al., 2018). A CellLight^®^ Reagent based on a GALNT2 (Golgi-resident enzyme) plasmid for labeling of Golgi apparatus was used according to the manufacturer’s instructions (C10592, Invitrogen, USA). The reagent was added to HCs at 5×10^6^ particles/ml at the time of medium change after attachment for 24 h incubation. GRASP55-mCherry and GRASP55-EGFP plasmids were used as previously described (Xiang and Wang, 2010; Zhang et al., 2018c). GalT-mCherry (87327, Addgene, USA) and GalT-EGFP (11929, Addgene, USA) plasmids were purchased. HCs were transfected with GRASP and GalT plasmids at 1 μg/ml using the Lipofectamine™ LTX Reagent with PLUS™ Reagent (15338-100, Invitrogen, USA) at the time of medium change for 24 h incubation. siRNAs for mouse VAMP3 (sc-41339, Santa Cruz Biotechnology, USA) and VAMP4 (sc-61767, Santa Cruz Biotechnology, USA) were purchased, with non-targeting siRNA being used as the negative control. MSCs were incubated in Opti-MEM for 6 h (31985-070, Invitrogen, USA) and were transfected with siRNAs at 50 nM for 48 h using the Lipofectamine™ RNAiMAX Reagent (13778-100, Invitrogen, USA).

#### GalNAc and GalNAc-conjugated siRNA experiments

For *in vivo* experiments, GalNAc (HY-33212, MedChemExpress, China) was injected subcutaneously at 500 mg/kg once per week. GalNAc-conjugated siRNAs were obtained from Alnylam Pharmaceuticals, USA, as 10 mg/ml stocks for G-si*Luc* and G-si*Anln*, as previously reported (Zhang et al., 2018a; Zhang et al., 2018b). The working concentrations of the siRNAs were 4 mg/ml diluted in PBS, and they were subcutaneously injected at 4 mg/kg once per week.

#### PHx surgeries

The PHx procedure was performed under sterile conditions by methods described previously (Hori et al., 2011; Mitchell and Willenbring, 2008). Isoflurane inhalation (R500IP, RWD Life Science, USA) was used to anesthetize the animals. For 70% PHx, the median and left lobes of liver were resected. For 80% MHx, the median, left and right posterior lobes of the liver were resected. Sham operation was performed by laparotomy.

#### ALF modeling and therapy

The mouse ALF model was established by APAP treatment, as stated previously (Barbier-Torres et al., 2017). Mice were fasted overnight, and APAP (A3035, Sigma-Aldrich, USA) was injected intraperitoneally at a lethal dose of 1 g/kg. NAC (A7250, Sigma-Aldrich, USA) was also injected intraperitoneally at 1 g/kg at 8 h post to APAP challenge. Control mice were injected with an equal amount of PBS.

#### Parabiosis

The parabiosis mouse model was established as previously stated (Liu et al., 2018). The mice to be joined in parabiosis were anaesthetized and shaved along the opposite lateral flanks. Incisions were made on the corresponding lateral aspects from the olecranon to the knee joint of each mouse. The olecranon and knee joints were each attached by sutures, and the dorsal and ventral skins were sewed together with continuous suture. Each pair of parabiotic mice was kept in a single cage after the operation.

#### Gross analyses

Body weights of mice were recorded weekly and food intake was recorded daily. Liver weights were recorded after sacrifice and gross morphology was characterized. Fasting experiments were performed after a fasting of 16 h.

#### Cage wheel running

As previously reported (Pistilli et al., 2011), spontaneous activity of mice was monitored using a cage running wheel system (Columbus Instruments, USA). Computer software (RMCWin, Delta Computer Systems, USA) specific to the equipment was used to quantify the total number of wheel revolutions. Data were expressed as distance covered per hour.

#### PET scanning and quantification of radiation intensity

PET scans were performed using an Inveon microPET scanner (Siemens Medical Solutions, Germany), as we previously documented (Sun et al., 2017). Briefly, mice were anesthetized using isoflurane at indicated time points after intravenous injection of ^64^Cu-labeled apoEVs and were scanned under PET. Reconstruction of PET images was done without correction for attenuation or scatter using a 3D ordered subsets expectation maximization algorithm. Image analysis was performed using ASI Pro VMTM software (Siemens Medical Solutions, Germany). At 48 h, mice were sacrificed and multiple organs/tissues were dissected and weighed for biodistribution analysis. The radiation intensity of ^64^Cu signals was measured in a well gamma-counter (Wallach Wizard, PerkinElmer, USA).

#### Liver histology and IF staining

At sacrifice, liver tissues were rapidly isolated and fixed overnight with 4% paraformaldehyde (PFA) (150146, MP Biomedicals, USA). For histological analyses, samples were dehydrated and embedded in paraffin, and 5 μm serial sections were prepared (RM2125, Leica, Germany). Sections then underwent either H&E staining (3801698, Leica, Germany), Masson’s trichrome staining (HT15-1KT, Sigma-Aldrich, USA) or TUNEL (G3250, Promega, USA) staining. For liver damage assessment, H&E staining sections were scored from 0-4 for sinusoidal congestion, vacuolization of the HC cytoplasm and parenchymal necrosis, according to Suzuki’s classification (Suzuki et al., 1993). Liver inflammation was scored from 0-3 indicating absent, mild, moderate or severe mononuclear and polymorphonuclear infiltrations (Sancho-Bru et al., 2007).

For IF staining, liver tissues were rapidly isolated, fixed in 4% PFA, cryoprotected with 30% sucrose, and embedded in optimal cutting temperature (OCT) compound (4583, Sakura Finetek, USA). The specimens were snap-frozen and sectioned into 10 μm sagittal sections (CM1950, Leica, Germany). For F-actin staining of HC borders, as we previously stated (Wang et al., 2014), sections were probed with phalloidin conjugated to either AlexaFluor-488 or Rhodamine (R37112/R37110, Invitrogen, USA) according to the manufacturer’s instructions, and counterstained with DAPI (ab104139, Abcam, UK). Percentages of binucleated and mononucleated HCs were quantified using ImageJ 1.47 software (National Institute of Health, USA) from at least four microscopic fields. For staining of Golgi apparatus or Kupffer cells, sections were treated with 0.3% Triton X-100 (X100-100ML, Sigma-Aldrich, USA) diluted in PBS for 20 min at room temperature, blocked with 5% BSA (700-100P, Gemini Bio, USA) dissolved in PBS for 1 h at room temperature, and stained with a rabbit anti-mouse Golgin84/Golga5 primary antibody (NBP1-83352, Novus Biologicals, USA) or a rabbit anti-mouse F4/80 primary antibody (ab6640, Abcam, UK) overnight at 4°C at a concentration of 1:100. After washing with PBS, sections were then stained with an Alexa Fluor 488-conjugated goat anti-rabbit secondary antibody (A-11008, Invitrogen, USA) for 1 h at room temperature at a concentration of 1:200, and counterstained with DAPI (ab104139, Abcam, UK).

#### Serum aminotransferase analyses and enzyme-linked immunosorbent assay (ELISA)

At sacrifice, whole peripheral blood was extracted from the mouse retro-orbital venous plexus, and serum was isolated by centrifugation at 3,000 g for 15 min at 4°C followed by centrifugation at 12,000 g for 15 min at 4°C. Concentrations of ALT and AST were determined using general chemistry kits (A7526-150 and A7561-150, Pointe Scientific, USA) according to the manufacturer’s instructions. Concentrations of tumor necrosis factor-alpha (TNF-α) and interleukin-6 (IL-6) were determined using ELISA kits (430907 and 431307, BioLegend, USA) according to the manufacturer’s instructions.

#### Ploidy analyses by PI staining

For detection of ploidy (Zhang et al., 2018b), freshly isolated primary HCs or cultured HCs collected after ploidy transition were fixed at 2×10^6^/ml in 75% ethanol overnight at −20°C.Cells were then incubated with 500 μl of PI/Rnase Staining Buffer (550825, BD Pharmingen, USA) at room temperature for 15 min and analyzed by flow cytometry.

#### IF staining of cells and apoEVs

For IF staining of cytoskeleton and organelles (Joseph et al., 2008; Kolobova et al., 2017), cultured HCs were fixed at 37°C for 15 min with 4% PFA dissolved in the microtubule stabilizing buffer (MTSB), which was composed of 50 mM piperazine-1,4-bisethanesulfonic acid (PIPES) (190257, MP Biomedicals, USA), 5 mM EGTA (E3889, Sigma-Aldrich, USA) and 5 mM MgSO_4_ (13142-100G, Honeywell, USA) in distilled water (pH = 7.0). After washing with MTSB, cells were permeabilized with ice-cold methanol for 10 min, washed with MTSB, and blocked with 5% BSA (700-100P, Gemini Bio, USA) in MTSB for 1 h at room temperature. Cells were then stained with a mouse anti-mouse GM130 primary antibody (610822, BD Transduction Laboratories, USA) alone or together with a rat anti-mouse α-tubulin primary antibody (MA1-80017, Invitrogen, USA) or a rabbit anti-mouse ac-α-tubulin (acetyl K40) primary antibody (ab179484, Abcam, UK) overnight at 4°C at a concentration of 1:100 in MTSB. After washing with MTSB, cells were then stained with Alexa Fluor 488-conjugated goat anti-mouse secondary antibody (A-11001, Invitrogen, USA) alone or together with Alexa Fluor 647-conjugated goat anti-rat secondary antibody (A-21247, Invitrogen, USA) or Alexa Fluor 647-conjugated goat anti-rabbit secondary antibody (A-21245, Invitrogen, USA) for 1 h at room temperature at a concentration of 1:200 in MTSB, and counterstained by DAPI (ab104139, Abcam, UK).

For IF staining of surface markers of apoEVs, after isolation, apoEVs were stained with rabbit anti-human/mouse VAMP3 primary antibody (13640, Cell Signaling Technology, USA), rabbit anti-mouse VAMP4 primary antibody (PA1-768, Invitrogen, USA), mouse anti-mouse GS15 primary antibody (610960, BD Transduction Laboratories, UK), rabbit anti-mouse C1q primary antibody (ab71940, Abcam, UK), or mouse anti-mouse TSP1 primary antibody (sc-59887, Santa Cruz Biotechnology, USA) at 4°C for 1 h at a concentration of 1:100 in PBS. After centrifugation and washing with PBS, apoEVs were then stained with FITC-conjugated goat anti-mouse secondary antibody (405305, BioLegend, USA) or FITC-conjugated donkey anti-rabbit secondary antibody (406403, BioLegend, USA) for 1 h at 4°C at a concentration of 1:200 in PBS, and counterstained with CellMask™ Deep Red (C10046, Invitrogen, USA).

#### Annexin V and lectin binding assay

Exposure of PtdSer on apoEVs was detected by FITC-conjugated Annexin V (640906, BioLegend, USA) binding at a concentration of 1:100 according to the manufacturer’s instructions and counterstained with CellMask™ Deep Red (C10046, Invitrogen, USA). Exposure of Gal or GalNAc on apoEVs, Exos and MSCs was respectively detected by FITC-conjugated peanut lectin (L7381, Sigma-Aldrich, USA) or Alexa Fluor 488-conjugated soybean lectin (L11272, Invitrogen, USA) binding both at a concentration of 1:100 for 1 h at 4°C, followed by counterstaining with CellMask™ Deep Red for membrane labeling (C10046, Invitrogen, USA). Fluorescence intensity was determined using a microplate reader (Synergy H1, Bio-Tek, USA) with Gen5 software (Bio-Tek, USA) at 488/517 nm for lectins and 649/666 nm for CellMask™ Deep Red.

#### Image acquisition and analyses

Bright-field images of cell morphology, liver H&E, Masson’s and TUNEL staining, and fluorescent images of liver engraftment of apoEVs, apoEV uptake *in vitro* and marker expression were obtained using an inverted microscope (Axio Observer 5, Zeiss, Germany). Fluorescent images of liver ploidy, apoEV uptake *in vivo,* Golgi and microtubule analyses were obtained using confocal microscopes (SP5-II, Leica, Germany; or LSM 900, Zeiss, Germany). Z-stacks of images were scanned at high resolution and were processed and reconstructed in 3 dimensions. Volocity software (PerkinElmer, USA) was applied for 3D analysis of apoEV interactions with the Golgi. Quantification of images was carried out with the ImageJ 1.47 software (National Institutes of Health, USA).

#### Flow cytometry

Sorting of HCs was performed using the MoFlo Astrios EQ Cell Sorter (Beckman Coulter, USA) with a 150 μm nozzle. Doublets were eliminated on the basis of pulse width, and ploidy populations were identified by Hoechst 33342-stained DNA contents using an ultraviolet 355 nm laser and a 425-440 nm bandpass filter. Ploidy analyses of HCs after PI staining were evaluated using a FACSCalibur Flow Cytometer (BD Biosciences, USA). Analyses of apoEV uptake efficacy, cellular apoptotic rates and apoEV marker expression were performed using a NovoCyte Flow Cytometer (ACEA Biosciences, USA). CellMask™ Deep Red staining was used to eliminate background noise signals when analyzing apoEVs. Flow cytometric data were processed using FlowJo v10 software (FlowJo LLC, USA).

#### Golgi isolation

Golgi apparatus was isolated from mouse liver or cultured HCs by sequential sucrose gradients using a Golgi Isolation Kit (GL0010-1KT, Sigma-Aldrich, USA), based on the protocol we previously established (Tang and Wang, 2015). Isolated Golgi proteins were extracted and the protein mass was determined using the BCA method (23225, Thermo Scientific, USA).

#### Transmission electron microscopy (TEM) analyses

Cultured HCs were collected by trypsin (12605-010, Invitrogen, USA) after 96 h in the ploidy transition protocol. HCs were then fixed with 2.5% glutaraldehyde (G5882, Sigma-Aldrich, USA) and 2.0% PFA (150146, Sigma-Aldrich, USA) solution in 0.1 M sodium cacodylate buffer (pH =7.4) (C0250, Sigma-Aldrich, USA) overnight at 4°C. After subsequent washes, the samples were post-fixed in 2.0% osmium tetroxide (75633-2ML, Sigma-Aldrich, USA) for 1 h at room temperature, and rinsed in distilled water prior to *en bloc* staining with 2% uranyl acetate (22400, Electron Microscopy Sciences, USA). After dehydration through a graded ethanol series, the specimens were infiltrated and embedded in the EMbed-812 resin (14120, Electron Microscopy Sciences, USA). Thin sections were stained with uranyl acetate and lead citrate (22410, Electron Microscopy Sciences, USA), and examined with a JEM-1010 electron microscope (JEOL, Japan) fitted with a digital camera (Hamamatsu Photonics, Japan) and image capture software (AMT Imaging, USA). Interphase 4N-HCs containing a binucleated profile with an intact nuclear envelope were selected to quantify distinguishable Golgi stacks per cell. Golgi stacks were identified using morphological criteria and were quantified using standard stereological techniques, as we previously stated (Souter et al., 1993; Xiang and Wang, 2010).

TEM analyses of apoEVs and liver tissue EVs were performed on collected pellets diluted in PBS. Diluted EVs were fixed with 1% glutaraldehyde (G5882, Sigma-Aldrich, USA) for 30 min at 4°C, absorbed onto glow-discharged 300-mesh heavy-duty carbon-coated formvar copper grids (ProSciTech, Australia) for 5 min, and the excess liquid was blotted on filter papers. Grids were washed twice with distilled water and negatively stained with 2.5% uranyl acetate (22400, Electron Microscopy Sciences, USA). Wide-field images encompassing multiple vesicles were taken on a JEM-1200EX electron microscope (JEOL, Japan).

#### NTA analyses

The size distribution of apoEVs and liver tissue EVs was measured using NanoSight NS300 (Malvern Panalytical, UK) and analyzed with NTA software (Malvern Panalytical, UK). Quantification of liver tissue EVs was also performed using ZetaView instrument S/N 19-447 (Particle Metrix, Germany) and analyzed with ZetaView analysis software (Particle Metrix, Germany).

#### RNA extraction and quantitative real-time polymerase chain reaction (qRT-PCR)

Total RNA was isolated from freshly isolated or cultured cells using the Aurum Total RNA Mini Kit (732-6820, Bio-Rad, USA) according to the manufacturer’s instructions. cDNA was synthesized using the Reverse Transcription Kit (4368814, Applied Biosystems, USA). qRT-PCR was performed using SYBR Green Master Mix (B21202, Bimake, USA) on the Real-Time PCR Systems (CFX96, Bio-Rad, USA; or LightCycler 96, Roche Life Science, Swiss). The primers used in this study are listed in Table S2. β-actin was used as the internal control.

#### Western blot assay

Western blot assay was performed on cultured MSCs or HCs, isolated apoEVs or liver tissue EVs, and isolated Golgi after lysis in the RIPA Lysis Buffer System with protease and phosphatase inhibitors (sc-24948, Santa Cruz Biotechnology, USA). Protein levels were quantified using the Pierce™ BCA Protein Assay Kit (23225, Thermo Scientific, USA). A total of 20 μg protein was separated in the 4%-12% NuPAGE gel (NP0321BOX/NP0322BOX, Invitrogen, USA) and was transferred to 0.2 μm nitrocellulose membranes (Millipore, USA). The membranes were then blocked with 5% BSA (700-100P, Gemini Bio, USA) at room temperature for 1 h, followed by incubation overnight at 4°C with the following primary antibodies: antibodies to GRASP55 (sc-271840), TGN38 (sc-166594), CD9 (sc-13118), CD63 (sc-5275), CD81 (sc-70803), INCENP (sc-376514), RacGAP1 (sc-166477) and RhoA (sc-418) were purchased from Santa Cruz Biotechnology, USA, and were used at concentrations of 1:200; antibodies to Stx5 (14151), Caspase 3 (9662), Cleaved caspase-3 (9664), VAMP3 (13640) and GAPDH (5174) were obtained from Cell Signaling Technology, USA, and were used at concentrations of 1:1000; the antibody to GM130 (610822) was purchased from BD Transduction Laboratories, USA, and was used at a concentration of 1:1000; the antibody to Golgin84 (NBP1-83352) was purchased from Novus Biologicals, USA, and was used at a concentration of 1:1000; antibodies to α-tubulin (MA1-80017) and VAMP4 (PA1-768) were purchased from Invitrogen, USA, and were used at concentrations of 1:1000; antibodies to ac-α-tubulin (ab179484), Lamin B1 (ab133741) and hVAMP3 (ab200657) were purchased from Abcam, UK, and were used at concentrations of 1:1000; the antibody to MKLP1/KIF23 (DF2573) was purchased from Affinity Biosciences, China, and was used at a concentration of 1:500; antibodies to GFP (SAB2702197) and β-actin (A5441) were purchased from Sigma-Aldrich, USA, and were used at concentrations of 1:1000. The membranes were then washed and incubated for 1 h at room temperature with horseradish peroxidase (HRP)-conjugated secondary antibodies (sc-516102/sc-2357, Santa Cruz Biotechnology, USA; or 7077, Cell Signaling Technology, USA). Immunoreactive proteins were detected using the SuperSignal™ West Pico PLUS Chemiluminescent Substrate (34580, Thermo Scientific, USA), the SuperSignal™ West Femto Maximum Sensitivity Substrate (34095, Thermo Scientific, USA), and the Autoradiography Film (LabScientific, USA) or the ChemiDoc MP Imaging System (Bio-Rad, USA).

#### Oxygen consumption assessment

Mitochondrial OXPHOS activity was measured using Mito-ID^®^ O_2_ Extracellular Sensor Kit (ENZ-51044, Enzo Life Sciences, USA) according to the manufacturer’s protocol, in which oxygen consumption was detected based on a fluorescent probe. Briefly, 10 μL probe was added to cultured HCs, and the fluorescence intensity was read immediately using the microplate reader (Synergy H1,Bio-Tek, USA) with the Gen5 software (Bio-Tek, USA) at 380/650 nm kinetically for 90 min, at an interval of 1 min. The mean value of fluorescence intensity of 3 wells per group was determined for the oxygen consumption curves (Lv et al., 2018).

#### ROS determination

Total intracellular ROS contents of cultured HCs were measured using the fluorescent probe 2’,7’-dichlorofluorescin diacetate (DCFDA) (ab113851, Abcam, UK) (Lv et al., 2018). Briefly, 25 mM DCFDA was added to cells and was incubated at 37°C for 30 min. Fluorescence intensity was determined using a microplate reader (Synergy H1, Bio-Tek, USA) with the Gen5 software (Bio-Tek, USA) with an excitation at 488 nm.

### QUANTIFICATION AND STATISTICAL ANALYSIS

Data are represented as the mean ± standard deviation (SD) unless otherwise indicated. Statistical significance was evaluated by two-tailed Student’s *t* test for two-group comparison, or by one-way analysis of variation (ANOVA) followed by Newman-Keuls post-hoc tests for multiple comparisons using Prism 5.01 software (GraphPad, USA). Log-rank tests were used for Kaplan-Meier survival curve comparisons. Values of *P* < 0.05 were considered statistically significant.

### MATERIALS AVAILABILITY

This study did not generate new unique reagents.

## DATA AND CODE AVAILABILITY

This study did not generate/analyze [datasets/code].

## Supplementary figure legends

**Figure S1. Identification of extracellular vesicles (EVs) and apoptosis-deficient mouse models used in this study. (A)** Schematic diagram demonstrating protocol for isolation of staurosporine (STS)-induced apoptotic extracellular vesicles (apoEVs). **(B)** Representative bright field images showing morphologies of cultured mesenchymal stem cells (MSCs) without or with STS treatment. Ctrl, control. Scale bars = 50 μm. **(C)** Representative fluorescent images showing terminal deoxynucleotidyl transferase dUTP nick end labeling (TUNEL, green) staining of apoptotic MSCs counterstained with DAPI (blue). Scale bars = 50 μm. **(D)** BCA analysis of the protein yield of apoEVs and exosomes (Exos) from 10^6^ MSCs. *N* = 3 per group. **(E)** Representative transmission electron microscopy (TEM) image showing the morphologies of apoEVs. Scale bar = 100 nm. **(F)** Nanosight analysis showing diameters of apoEVs. (**G-I**) Representative fluorescent images show apoEV (labeled by CellMask™ Deep Red, red) surface exposure of phosphatidylserine (PtdSer, Annexin V labeling) and expression of complement component 1q (C1q) and thrombospondin 1 (TSP1) (green). Flow cytometric analysis determined percentages of positively stained apoEVs. Scale bars = 25 μm. **(J)** Western blot showing marker expression in apoEVs and source MSCs. **(K)** Flow cytometric analysis showed apoptosis of peripheral blood mononucleated cells (PBMNCs). PBMNCs from wild-type (WT), *Fas* mutant (*Fas^mut^*) or *Caspase-3* knockout (*Casp3^-/-^*) mice, staining with Annexin V and 7AAD. Percentages of Annexin V^+^ cells were quantified as apoptotic rates. ND, not detected. *N* = 3 per group. **(L)** Flow cytometric analysis showed apoptosis of bone marrow mononucleated cells (BMMNCs). BMMNCs from WT, *Fas^mut^* or *Casp3^-/-^* mice were stained with Annexin V and 7AAD. Percentages of Annexin V^+^ cells were quantified as apoptotic rates. *N* = 3 per group. **(M)** Flow cytometric analysis showed apoptosis of liver non-parenchymal cells (NPCs). NPCs from WT, *Fas^mut^* or *Casp3^-/-^* mice were stained with Annexin V and 7AAD. Percentages of Annexin V^+^ cells were quantified as apoptotic rates. *N* = 3 per group. **(N)** Schematic diagram demonstrating the liver cell and EV isolation protocol. HCs, hepatocytes. **(O)** Representative TEM image showing the morphologies of liver EVs. Scale bar = 50 nm. **(P)** Nanosight analysis showing diameters of liver EVs. (**Q**) Western blot showing marker expression in liver EVs and HCs. Data represent mean ± standard deviation. ***, *P* < 0.0001. Statistical analyses were performed by one-way analysis of variance followed by the Newman-Keuls post hoc tests.

**Figure S2. Phenotypic profiles of *Fas* mutant (*Fas^mut^*) mice after apoEV infusion. (A)** Gross view images of 16-week-old mice and quantification of body weight (BW). WT, wild type. Scale bars = 1 cm. *N* = 6 per group. ***, *P* < 0.0001, *Fas^mut^* mice compared to WT mice; NS, not significant, *P* > 0.05, *Fas^mut^* mice after apoEV infusion compared to *Fas^mut^* mice. **(B)** Quantification of food intake of 16-week-old mice. NS, not significant, *P* > 0.05. **(C)** Spontaneous activity of 16-week-old mice was assessed by running distance per hour using a cage running wheel system. Gray area indicates dark phase. *N* = 3 per group. **(D)** Enzyme-linked immunosorbent assay (ELISA) detected serum levels of tumor necrosis factor-alpha (TNF-α) and interleukin-6 (IL-6) in 16-week-old mice. NS, not significant, *P* > 0.05. *N* = 6 per group. **(E)** ZetaView analysis showing liver tissue extracellular vesicles (EVs) and corresponding quantification. NS, not significant, *P* > 0.05. *N* = 3 per group. **(F)** Masson’s trichrome staining showing liver tissue and corresponding quantification of fibrotic area percentages. Scale bars = 50 μm. NS, not significant, *P* > 0.05. *N* = 6 per group. **(G)** Hepatic inflammation scores were examined based on pathological parameters. NS, not significant, *P* > 0.05. *N* = 6 per group. **(H)** Tracing of PKH26-labeled (for membrane) or aggregation-induced emission luminogen (AIEgen, for mitochondria)-labeled apoEVs (red) in liver, with Kupffer cell membranes stained with F4/80 (green) and counterstaining with DAPI for DNA (blue). ApoEVs were intravenously injected and liver samples were collected at 24 h after the infusion. Representative confocal images showed macrophages. Scale bars = 5 μm. Data represent mean ± standard deviation. Statistical analyses were performed by one-way analysis of variance followed by the Newman-Keuls post hoc tests.

**Figure S3. *Caspase-3* knockout (*Casp3^-/-^*) mice demonstrate hepatic disorders rescued by apoEV infusion. (A)** Schematic diagram demonstrates the study design of apoEV infusion in *Casp3^-/-^* mice. ApoEVs were infused intravenously (*i.v.*) at a protein concentration of 5 μg/g. **(B)** ZetaView analysis showing liver tissue extracellular vesicles (EVs) and corresponding quantification. WT, wild type. *N* = 3 per group. **(C)** Flow cytometric analysis showing Annexin V^+^ liver tissue EVs (apoEVs) with corresponding quantification. *N* = 3 per group. **(D)** Hematoxylin and eosin (H&E) staining of liver tissues in periportal vein (PV) area. Hepatic injury scores were examined based on pathological parameters. Scale bars = 50 μm. *N* = 4 per group. **(E)** Serum alanine aminotransferase (ALT) levels were determined. *N* = 4 per group. **(F)** Representative liver fluorescent images show hepatocytes (HCs) with different nuclei (blue, DAPI for DNA) and their cell borders (green, phalloidin for F-actin). # indicates binucleated HCs. Percentages of binucleated HCs were accordingly quantified. Scale bars = 25 μm. *N* = 4 per group. **(G)** Diploid HCs (2N-HCs) were analyzed by flow cytometry after propidium iodide (PI) staining. *N* = 4 per group. **(H)** Tracing of PKH26-labeled apoEVs (red) in sorted tetraploid HCs (4N-HCs), with Golgi apparatus being demonstrated by a plasmid of *N*-acetylgalactosaminyltransferase 2 (GALNT2)-GFP (green) and counterstaining by DAPI for DNA (blue). Imaging analysis was performed to quantify HC percentages with Golgi fragmentation. Scale bars = 5 μm. *N* = 4 per group. **(I)** Golgi apparatus was isolated from cultured HCs and Golgi protein mass was determined by using the BCA method. *N* = 4 per group. Data represent mean ± standard deviation. *, *P* < 0.05; **, *P* < 0.01; ***, *P* < 0.0001; NS, not significant, *P* > 0.05. Statistical analyses were performed by one-way analysis of variance followed by the Newman-Keuls post hoc tests.

**Figure S4. Tetraploid hepatocytes (4N-HCs) are weaker in metabolic activity and more prone to oxidative stress than diploid hepatocytes (2N-HCs). (A)** Flow cytometric sorting of primary mouse HCs based on Hoechst 33342 staining. Representative bright field images showing morphologies of 2N- and 4N-HCs. Scale bars = 10 μm. **(B)** Oxidative phosphorylation (OXPHOS) metabolic activities of sorted HCs were analyzed by oxygen consumption kinetics. WT, wild type; *Fas^mut^*, *Fas* mutant; apoEVs, apoptotic extracellular vesicles. *N* = 3 per group. Curves are depicted with mean values. **(C)** Intensity of cellular reactive oxygen species (ROS) in sorted HCs were analyzed by the 2□,7□-dichlorofluorescin diacetate (DCFDA) probe. *N* = 3 per group. Data represent mean ± standard deviation. **(D)** Quantitative real-time polymerase chain reaction (qRT-PCR) showed mRNA expression of multiple antioxidants in sorted HCs normalized to *β-actin*. Nfe2l2, nuclear factor erythroid 2-related factor 2; Sod2, superoxide dismutase 2; Cat, catalase; Nqo1, NAD(P)H quinone dehydrogenase 1; Gclc, glutamate-cysteine ligase catalytic subunit; Gpx1, glutathione peroxidase 1; Gsr, glutathione reductase. *N* = 3 per group. Data represent mean ± standard deviation. **(E)** OXPHOS metabolic activities of unsorted primary HCs were analyzed based on oxygen consumption kinetics. *N* = 3 per group. Curves are depicted with mean values. **(F)** Intensity of cellular ROS in unsorted primary HCs was determined by the DCFDA probe. *N* = 3 per group. Data represent mean ± standard deviation. **, *P* < 0.01; ***, *P* < 0.0001. Statistical analyses were performed by Student’s *t* test for two-group analysis or one-way analysis of variance followed by the Newman-Keuls post hoc tests for multiple group comparisons.

**Figure S5. Circulating extracellular vesicles (EVs) in parabiosis mice contribute to liver homeostasis. (A)** Schematic diagram demonstrates the study design of parabiosis using *Fas* mutant (*Fas^mut^*) mice and green fluorescent protein (GFP) transgenic (*GFP^+/+^*) mice. *N*-acetylgalactosamine (GalNAc) was injected subcutaneously (*s.c.*) at 500 mg/kg once per week for inhibiting hepatocyte (HC) uptake of apoEVs. **(B)** Tracing of GFP-labeled particles (green) in liver of *Fas^mut^* mice, with HC cell borders being stained with phalloidin for F-actin (red) and counterstaining with DAPI for DNA (blue). Representative confocal images show binucleated HCs. Scale bar = 10 μm. **(C)** Hematoxylin and eosin (H&E) staining of liver tissues in periportal vein (PV) area. Hepatic injury scores were examined based on pathological parameters. Scale bars = 50 μm. *N* = 4 per group. **(D)** Serum alanine aminotransferase (ALT) levels were determined. *N* = 4 per group. **(E)** Representative liver fluorescent images show HCs with different numbers of nuclei (blue, DAPI for DNA) and their cell borders (green, phalloidin for F-actin). # indicates binucleated HCs. Percentages of binucleated HCs were accordingly quantified. Scale bars = 25 μm. *N* = 4 per group. **(F)** Diploid HCs (2N-HCs) were analyzed by flow cytometry after propidium iodide (PI) staining. *N* = 4 per group. **(G)** Representative Golgin84 immunofluorescent staining (green) images of liver Golgi counterstained with DAPI (blue). Imaging analysis was performed to quantify Golgi area percentages. Scale bars = 5 μm. *N* = 4 per group. **(H)** Golgi apparatus was isolated from liver and Golgi protein mass was determined using the BCA method. *N* = 4 per group. Data represent mean ± standard deviation. **, *P* < 0.01; ***, *P* < 0.0001. Statistical analyses were performed by one-way analysis of variance followed by the Newman-Keuls post hoc tests.

**Figure S6. ApoEVs are characterized by a general combined surface signature for liver regulation. (A-C)** Flow cytometric analyses were performed to determine percentages of positively stained apoEVs by Annexin V for surface exposure of phosphatidylserine (PtdSer), by FITC-conjugated lectins for surface exposure of galactose (Gal) and *N*-acetylgalactosamine (GalNAc), and by a specific antibody for surface expression of vesicle-associated membrane protein 3 (VAMP3). The general apoptotic vesicular signature (AVS) was identified with apoEVs from B-cell lymphoma-2 (Bcl-2) homology domain 3 (BH3)-mimetic ABT-737-induced apoptosis (*via* the intrinsic pathway) of human mesenchymal stem cells derived from the umbilical cord (hMSCs, **A**), anti-Fas-induced apoptosis (*via* the extrinsic pathway) of activated mouse T lymphocytes (**B**), and staurosporine (STS)-induced apoptosis of the mouse fibroblast cell line NIH/3T3 (**C**). (**D**) Graphical summary demonstrates the main findings of this study. ApoEVs were characterized by the specific AVS and engrafted in liver. ApoEVs are recognized and uptaken by hepatocytes (HCs) through the asialoglycoprotein receptor (ASGPR), in which they assemble with the Golgi apparatus to form a chimeric ApoEV-Golgi Complex (AGC) based on the soluble *N*-ethylmaleimide-sensitive factor attachment protein receptor (SNARE) mechanism. ApoEV uptake and AGC assembly activate downstream functional cascades including microtubule acetylation and cytokinesis of HCs, which are indispensable for ploidy transition of HCs to safeguard diploid (2N) rather than tetraploid (4N) HC populations. Ultimately, apoEVs are of functional significance for maintaining liver homeostasis, promoting liver regeneration and mediating therapy for acute liver failure.

**Supplementary tables** (*For Table S1, see the Key Resources Table*)

**Table S2.**
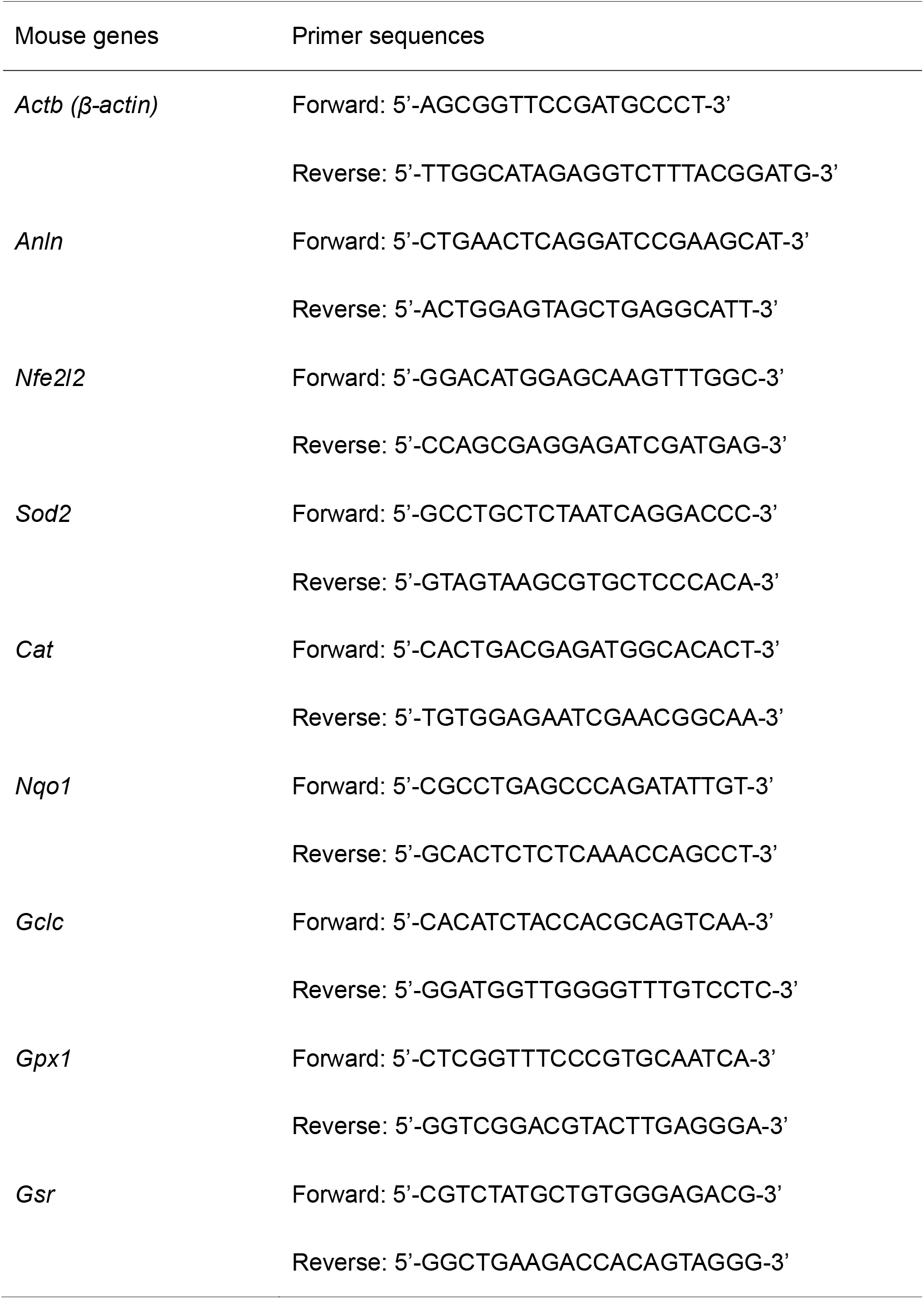
Primers used in this study.

