## Supplementary figures and images for "Apoptotic Extracellular Vesicles (ApoEVs) Safeguard Liver Homeostasis and Regeneration *via* Assembling an ApoEV-Golgi Organelle"

### Figure S1

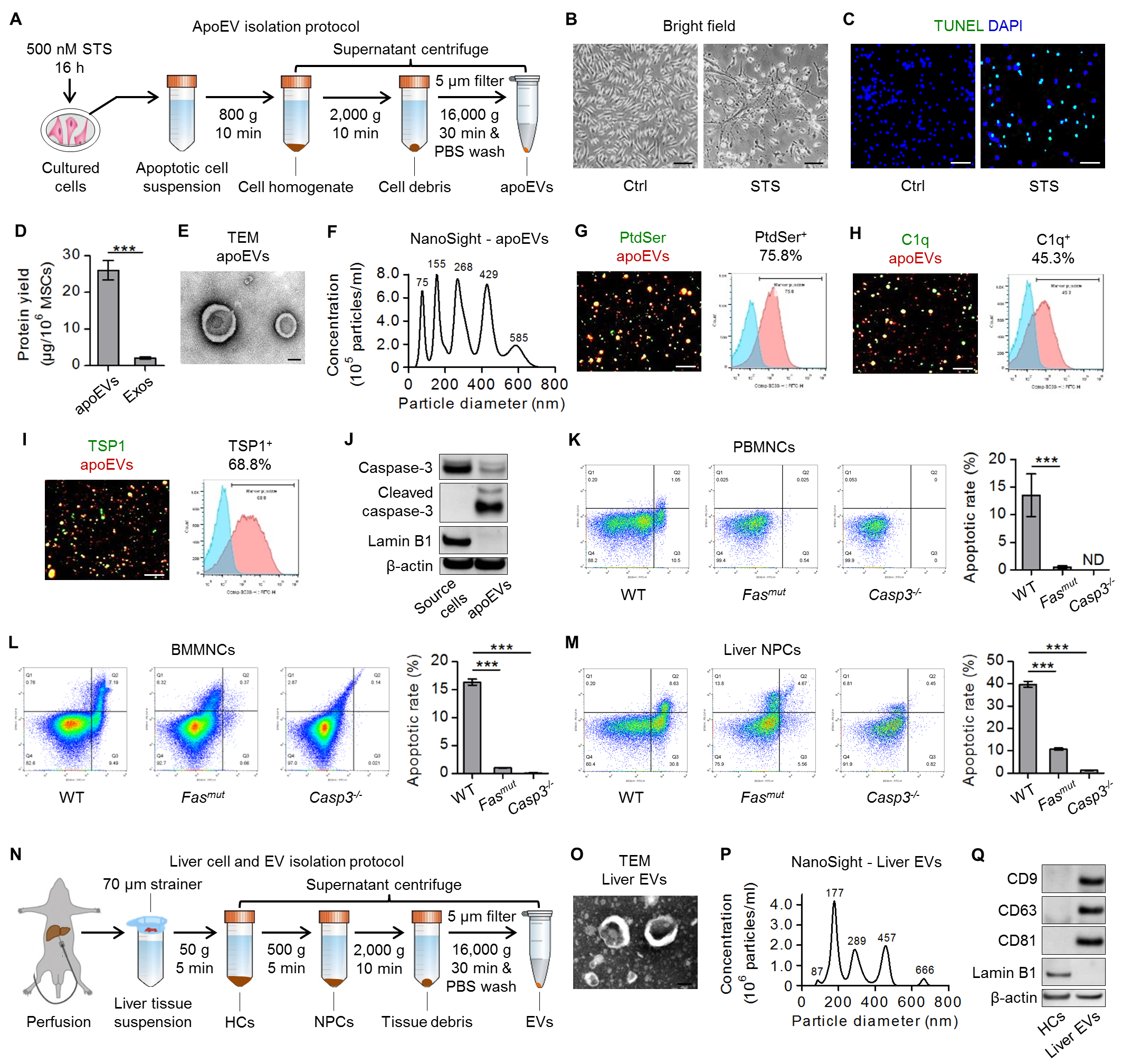

### Figure S2

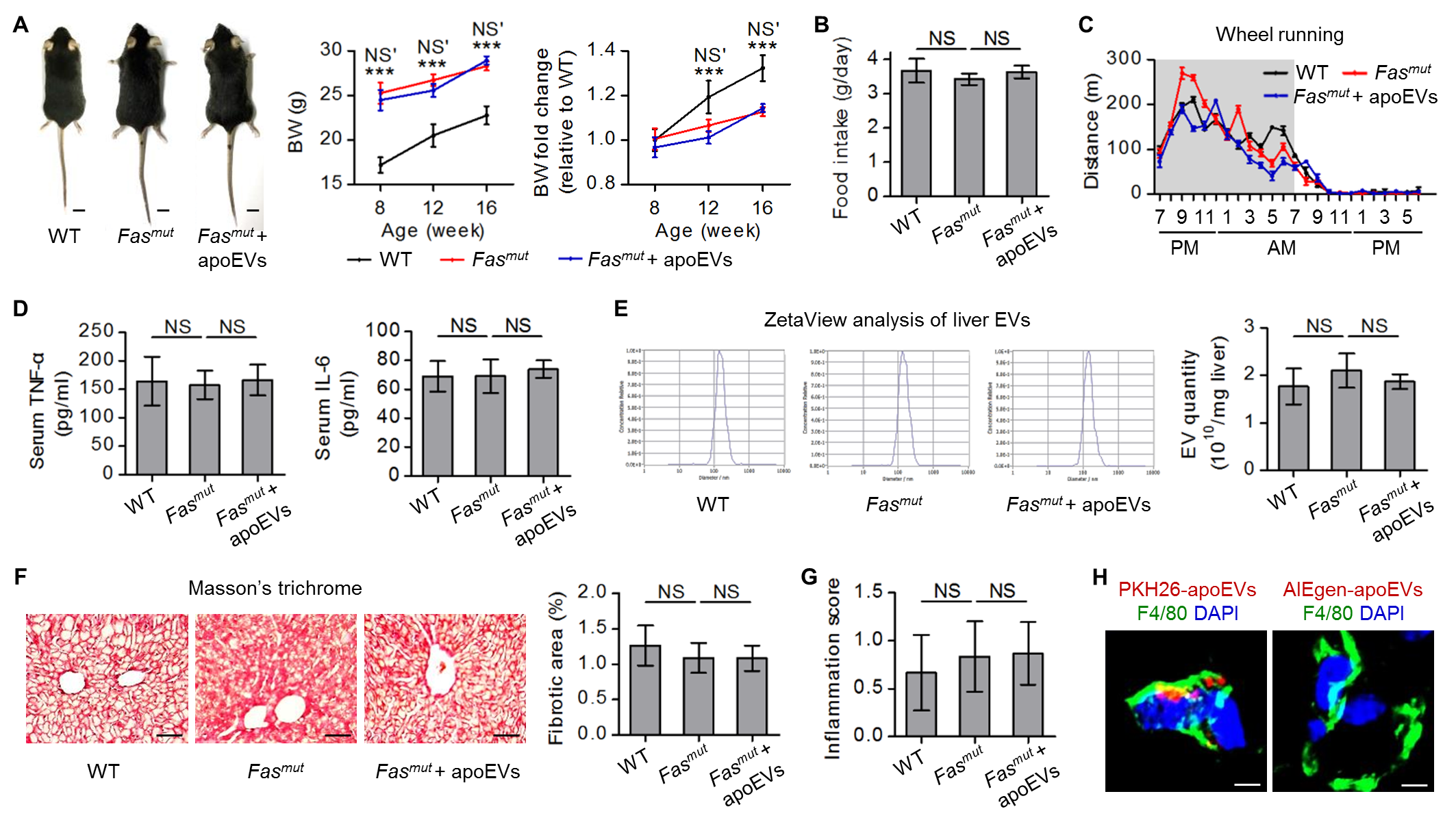

### Figure S3

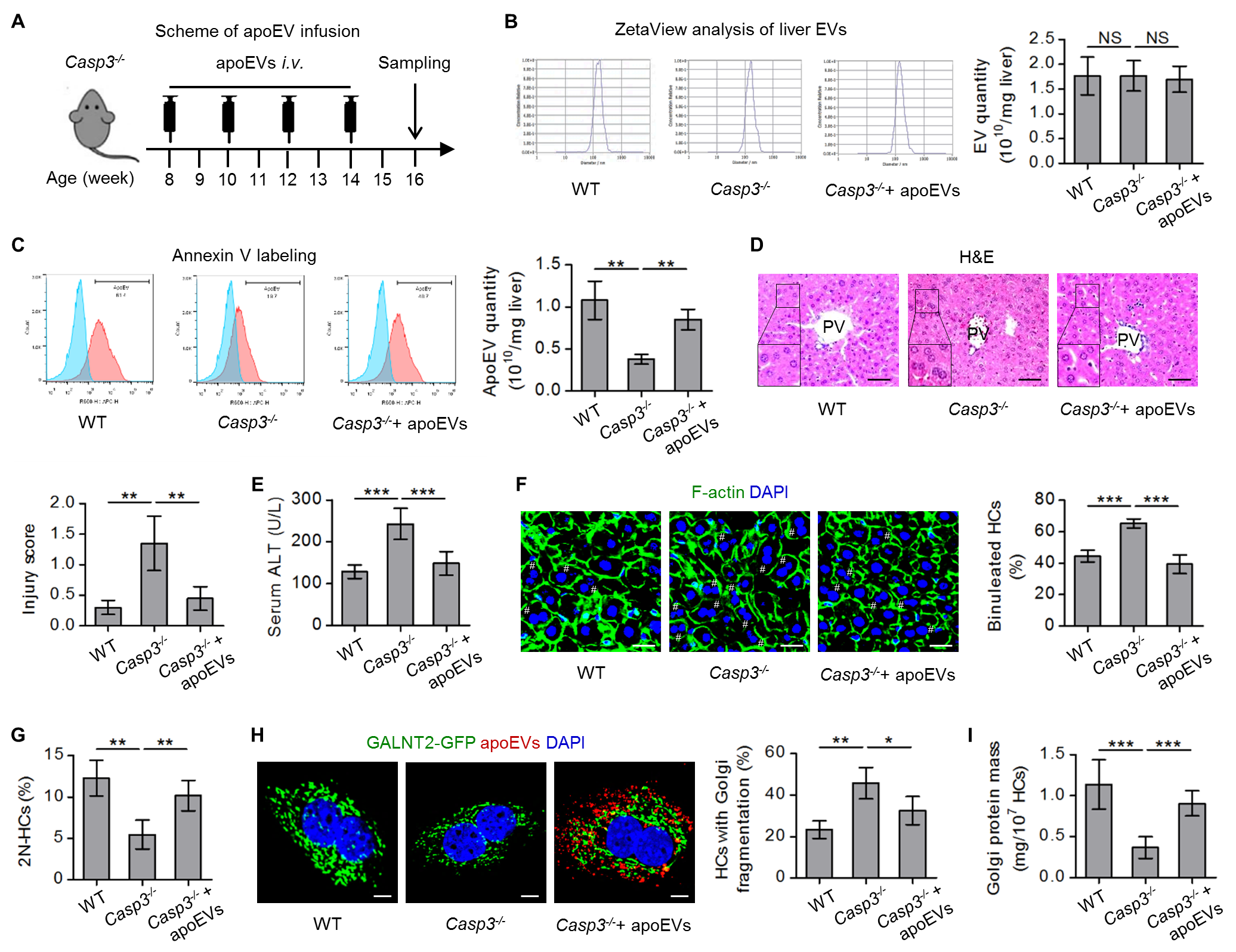

### Figure S4

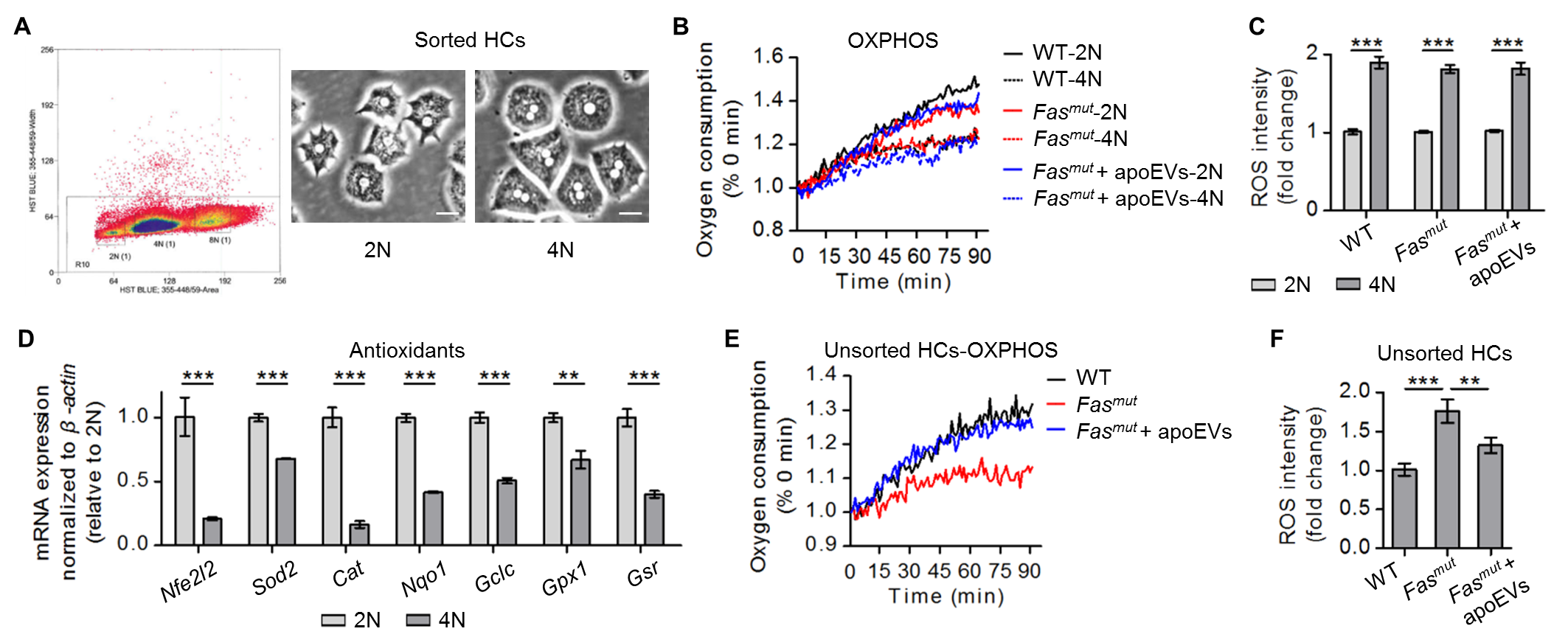

### Figure S5

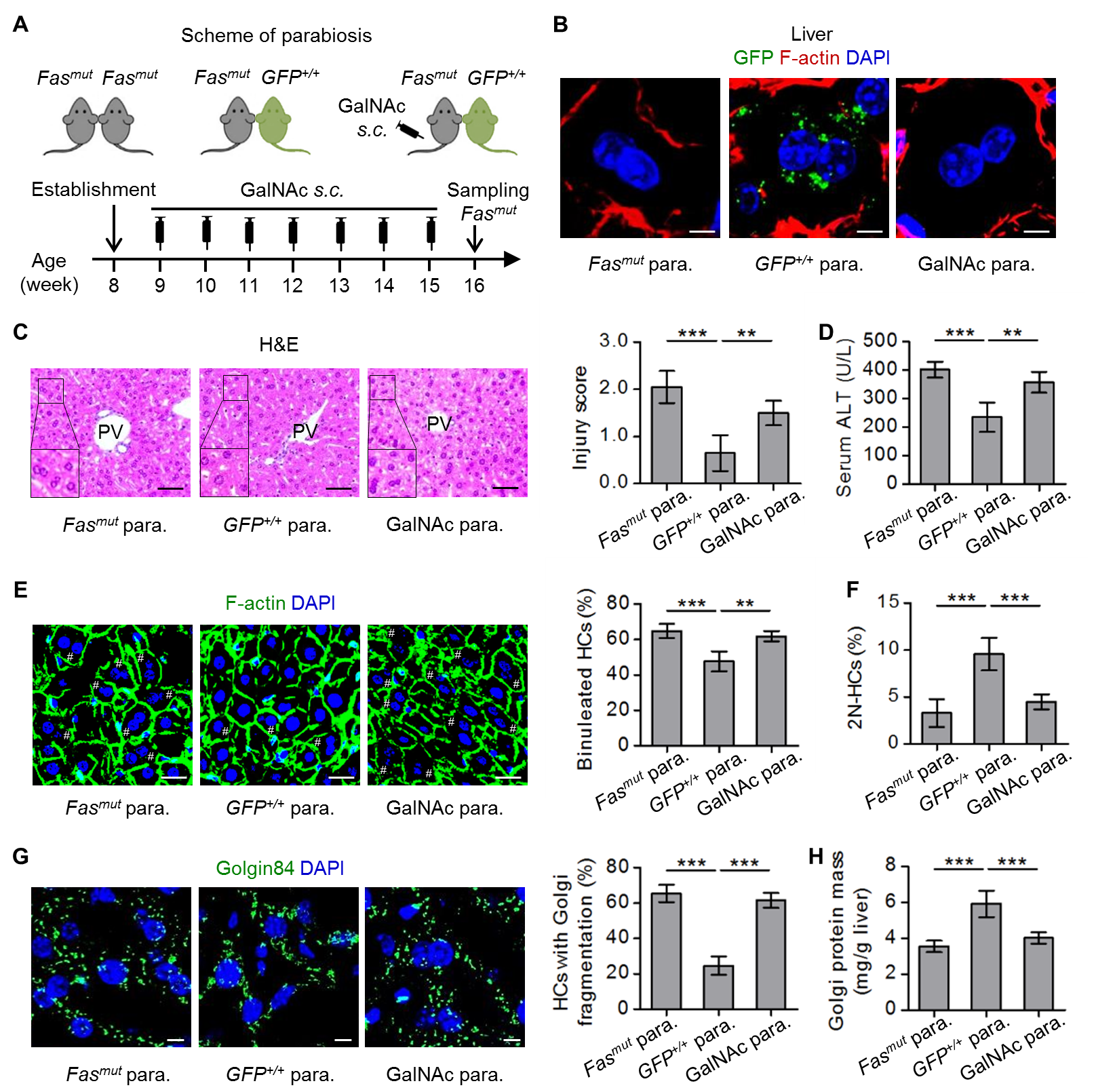

### Figure S6

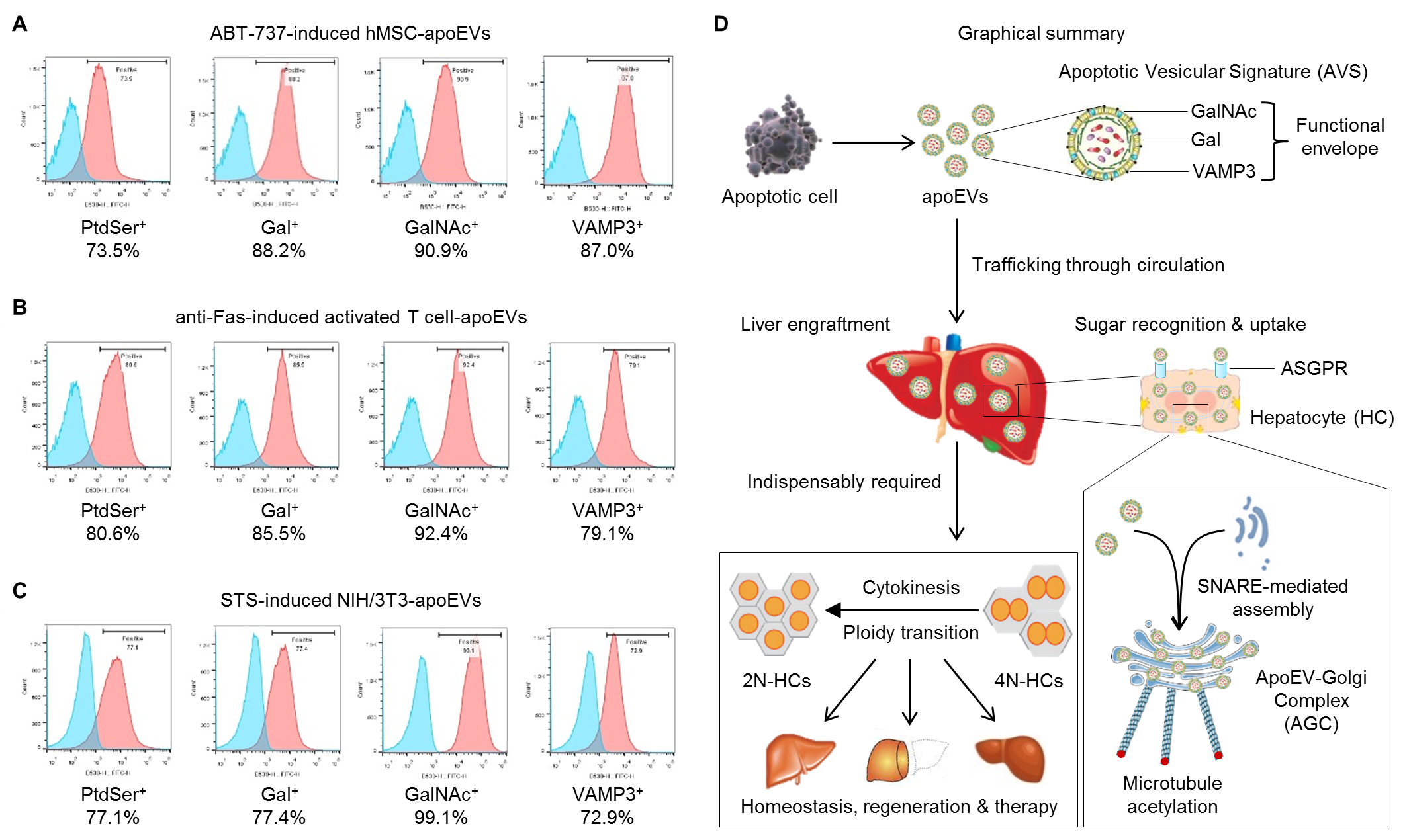
